# Co-developing climate services for public health: stakeholder needs and perceptions for the prevention and control of *Aedes*-transmitted diseases in the Caribbean

**DOI:** 10.1101/587188

**Authors:** Anna M. Stewart-Ibarra, Moory Romero, Avery Q. J. Hinds, Rachel Lowe, Roché Mahon, Cedric J. Van Meerbeeck, Leslie Rollock, Marquita Gittens-St. Hilaire, Sylvester St. Ville, Sadie J. Ryan, Adrian R. Trotman, Mercy J. Borbor-Cordova

## Abstract

**Background:** Small island developing states (SIDS) in the Caribbean region are challenged with managing the health outcomes of a changing climate. Health and climate sectors have partnered to co-develop climate services to improve the management of these diseases, for example, through the development of climate-driven early warning systems. The objective of this study was to identify health and climate stakeholder perceptions and needs in the Caribbean, with respect to the development of climate services for arboviruses (e.g. dengue, chikungunya, and Zika).

**Methods:** Stakeholders included public decision makers and practitioners from the climate and health sectors at the regional (Caribbean) level and from the countries of Dominica and Barbados. From April to June 2017, we conducted interviews (n=41), surveys (n=32), and national workshops with stakeholders. Survey responses were tabulated and audio recordings were transcribed and analyzed using qualitative coding to identify responses by research topic, country/region, and sector.

**Results:** Health practitioners indicated that their jurisdiction is currently experiencing an increased risk of diseases transmitted by *Ae. aegypti* due to climate variability, and most anticipated that this risk will increase in the future. National health sectors reported financial limitations and a lack of technical expertise in geographic information systems (GIS), statistics, and modeling, which constrained their ability to implement climate services for arboviruses. National climate sectors were constrained by a lack of personnel. Stakeholders highlighted the need to strengthen partnerships with the private sector, academia, and civil society. They identified a gap in local research on climate-arbovirus linkages, which constrained the ability of the health sector to make informed decisions. Strategies to strengthen the climate-health partnership included a top-down approach by engaging senior leadership, multi-lateral collaboration agreements, national committees on climate and health, and shared spaces of dialogue. Mechanisms for mainstreaming climate services for health operations to control arboviruses included climate-health bulletins and an online GIS platform that would allow for regional data sharing and the generation of spatiotemporal epidemic forecasts.

**Conclusions:** These findings support the creation of interdisciplinary and intersectoral communities of practices and the co-design of climate services for the Caribbean public health sector. By fostering the effective use of climate information within health policy, research and practice, nations will have greater capacity to adapt to a changing climate.

## Introduction

Small island developing states (SIDS) in the Caribbean region are highly susceptible to the health impacts of climate variability and long-term changes in climate [1,2]. Impacts include increased risk of communicable diseases, such as mosquito-borne arboviruses, and noncommunicable diseases, such as cardiovascular complications associated with heat stress. SIDS face similar challenges—small populations who are repeatedly exposure to extreme climate events (e.g., droughts and tropical storms), limited global political power, reliance on imported goods, and difficulty preventing and responding to disasters due to resource constraints [1,3,4]. Caribbean SIDS will likely experience more extreme climate events in the future due to climate change, increasing the social and economic burden of climate-sensitive health outcomes.

Dengue fever, chikungunya and Zika fever, arboviral diseases transmitted by the *Aedes aegypti* mosquito, are among the top public health concerns in the Caribbean region [5,6]. The Caribbean Public Health Agency (CARPHA) recently issued an advisory for the possibility of a severe dengue epidemic in 2019 [7], given a rise in dengue activity in Latin America, the increasing burden of arboviruses over the last number of years, and the large gap in time since the last dengue epidemic in the Caribbean region, which occurred in 2009. The Caribbean is the region of the Americas with the highest incidence of dengue [8]. Vector control is the main public health interventions to prevent and control disease outbreaks through insecticide application, elimination of larval habitat sites, public education and community mobilization [9]. Despite these efforts, the annual number of dengue cases in the region increased from an estimated 136,000 to 811,000 cases between 1990 and 2013, with case estimates adjusted to account for underreporting [8]. Novel tools and management strategies are urgently needed to increase the capacity of the public health sector to prevent and respond to arboviral disease outbreaks.

Changes in local climate can influence *Ae. aegypti* physiology and population dynamics, thereby affecting disease transmission. Warmer ambient temperatures increase the probability of arbovirus transmission by *Ae. aegypti*, with optimum transmission at 28.5°C; however, transmission is improbable in extreme heat (>34°C) [10]. Both excess rainfall and drought conditions can potentially increase mosquito densities, depending on the characteristics of the build environment and anthropogenic water storage [11,12]. In the Caribbean region, studies have documented the effects of climate on dengue and *Ae. aegypti* in Barbados [11,13–16], Cuba [17], Puerto Rico [18–20], Jamaica [16,21], Trinidad and Tobago [13,16], and Guadeloupe [22].

Given the linkages between arboviruses, *Ae. aegypti*, and climate, the World Health Organization and experts in the Caribbean have recommended developing climate-driven early warning systems (EWS) and models to forecast arbovirus outbreaks [23,24]. These tools are known as climate services -- tailored products for a specific sector that allow decision makers and practitioners to plan interventions. For example, an EWS for arboviruses could inform decisions about when and where to deploy public health interventions to prevent an epidemic in the context of an impending climate event [25]. A recent study by Lowe et al. [11] found that dengue transmission in Barbados increased one month after a particularly wet month and five months after a drought event was observed, and the model was able to accurately predict outbreak versus non-outbreak months. This study demonstrated the potential to develop an operational climate-driven forecast model to predict arbovirus outbreaks in the eastern Caribbean.

The perceptions, needs, and interests of stakeholders from the health and climate sectors should ideally drive the development and implementation of a forecast model through an iterative engagement process with modelers and other scientists [26,27]. The end-users of climate information are a diverse group of actors with distinct needs and interests[28,29]. Morss et al. [28] report insightful lessons learned as scientists who attempt to communicate flood risk to the public sector. They state, “Decision makers are not a coherent entity, but a collection of individuals, each of whom uses different information to address different goals in a unique context.” An arbovirus forecast should be developed with a realistic understanding of the present-day public health response capacity, which may be constrained by resources, information, prior experiences, and other actors or institutions [30]. To guide this process, the Global Framework for Climate Services (GFCS) was developed as the policy mechanism to support the development of climate services for the health sector and other key sectors [31]. The GFCS aims for stakeholder engagement between health and climate actors at all levels to promote the effective use of climate information within health research, policy and practice [31].

Prior studies of health sector perceptions of climate have focused on the perceptions of the health sector with respect to the impacts of long-term climate change and climate variability. However, few studies (primarily for malaria early warning systems, MEWS, e.g. [32–34]) address health sector needs and interests with respect to climate-driven epidemic forecasts. Studies from the United States and Canada have analyzed the perceptions and engagement of public health practitioners in the context of long-term climate change and impacts on overall health [35–39]. One study from China assessed health sector perceptions of dengue and climate change [40]. In the Caribbean, a study found that health providers perceived mosquito-borne disease as increasing due to changing seasonal patterns [41], whereas another study found that health practitioners had limited understanding of the effects of climate variability on health [42]. Climate practitioners in Jamaica were generally aware of the health implications of climate change for heat stress, respiratory diseases, and vector borne diseases [43].

The objective of this study was to identify health and climate stakeholder perceptions and needs in the Caribbean, with respect to the development of climate services for arboviruses. We addressed four key areas, based on the GFCS health exemplar goals [31]:

1. What are the perceptions of climate-health or climate-arbovirus linkages?
2. Who are the key actors engaged in climate-arbovirus surveillance and control, and how can communication and partnerships amongst these actors be strengthened?
3. What are the current capabilities of the health and climate sectors to implement a climate-driven arbovirus EWS, and what capacities need to be strengthened so that the health sector can effectively access, understand and use climate/weather information for decision-making?
4. What climate/weather data are currently used by the health sector for arbovirus control, what added value does it provide, and how can climate/weather data be effectively tailored for arbovirus control operations?

## Methods

### Ethical statement

The study protocols were reviewed and approved (or deemed exempt) by the Institutional Review Board (IRB) of the State University of New York Upstate Medical University, the IRB of the University of the West Indies, Cave Hill Campus on behalf of the Ministry of Health of Barbados, and the Ministry of Health and Environment of Dominica. No informed consent was required, as all participants were adults (>18 years of age), were public sector employees, and no identifying information was gathered.

### Study sites

This study focused on the perspectives of health and climate stakeholders from the countries of Barbados and Dominica (Figure 1), SIDS in the eastern Caribbean, as well as regional Caribbean stakeholders. There is a high burden of arboviral diseases in both countries [44–48] (Table 1). These countries were selected because of the regional and national interest in building on previous projects, wherein the health sector identified climate services as a top priority for the management of arboviruses.

**Figure 1:**
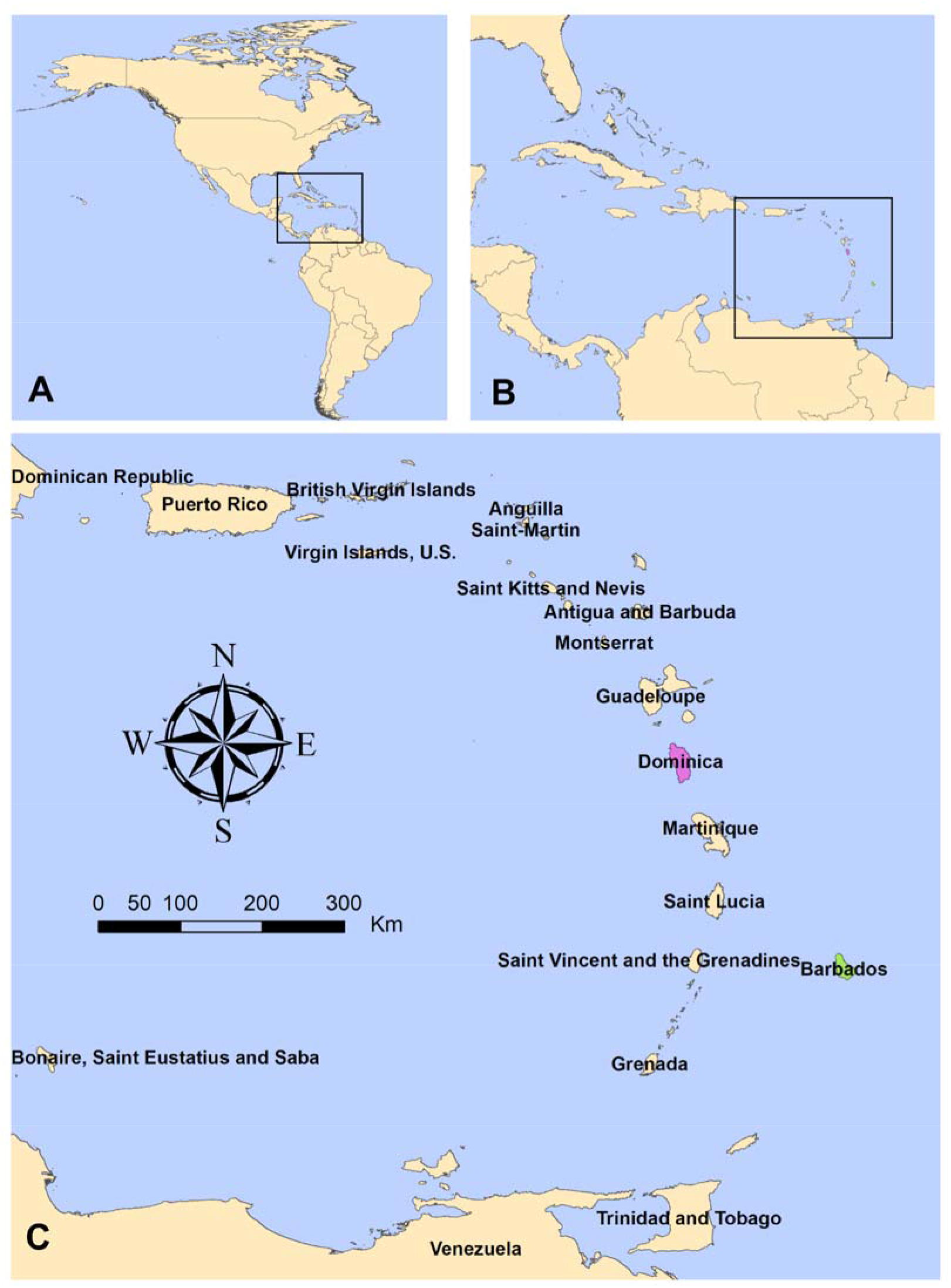
Map of the study region, showing A. The location of the Caribbean region within the Americas, with inset B. showing the archipelago of islands making up the Caribbean, and their location within Meso-America, with inset C. the location of Dominica (purple) and Barbados (green) within the islands in the region. This map was created using freely available country boundary data from GADM.org, rendered in ArcGIS, and image files created using GIMP freeware.

**Table 1.** Arboviral disease cases in Barbados and Dominica.

|  | Barbados | Dominica |
| --- | --- | --- |
| Mean annual dengue cases (2012-2016) [82]* | 2,274 | 169 |
| Mean annual dengue incidence (2012-2016) per 10,000 people [82]* | 80.0 | 23.5 |
| Total chikungunya cases since 2013 [83] | 1,833 suspected<br>114 confirmed | 3,590 suspected<br>173 confirmed |
| Total Zika cases since 2016 [46,47] | 705 suspected<br>150 confirmed | 1,263 suspected<br>79 confirmed |

\*Includes suspected and confirmed cases

Barbados (pop. 284,996; land area: 439 km^2^) has a service-based economy, with tourism accounting for 12% of the gross domestic produce (GDP). However, tourism is a water-intensive sector, and droughts threaten to reduce the already limited freshwater resources [49]. From 2011-2015, Barbados was selected as the country in the Western hemisphere for the WHO project on climate change adaptations strategies for human health, funded by the Global Environment Facility (GEF) Special Climate Change Fund (SCCF) [50]. The Ministry of Health is responsible for arbovirus and vector surveillance and control.

Dominica (pop. 73,543; land area: 750 km^2^) is characterized by abundant freshwater resources, forest and rugged terrain. Eco-tourism is becoming increasingly significant to its economy. Dominica was selected as the health exemplar for the GFCS, resulting in a national consultation on climate and health vulnerability in 2015-2016 that included vector borne diseases (VBDs), food safety, and water borne diseases [51]. The country was devastated by Hurricane Maria in September 2017, a category 5 hurricane that damaged 90% of buildings, resulting in USD 1.3 billion in damages, the equivalent of 224% of Dominica’s GDP in 2016 [52]. This study was conducted in the months prior to Hurricane Maria. The Ministry of Health and Environment is responsible for arbovirus and vector surveillance and control.

The national health agencies of both countries are supported by the Caribbean Public Health Agency (CARPHA) and the Pan American Health Organization (PAHO), the regional arm of the WHO. Each country has its own national meteorological and hydrological service (NMHS) supported by the Caribbean Institute for Meteorology and Hydrology, the technical arm of the Caribbean Meteorological Organization (CMO). Details regarding the mandates and capabilities of the public health and climate national and regional organizations, with respect to arbovirus and vector surveillance and control, and climate monitoring and forecasting, are provided in S1 Text.

### Surveys and interviews

We collected data from key stakeholders from the climate and health sectors spanning senior leadership, managers, and expert practitioners. Stakeholders from the health sector were engaged in arbovirus epidemiology and vector control, or environmental health, at national and regional (Caribbean) agencies. Stakeholders from the climate sector were individuals involved in the development of climate services for the Caribbean region and managers/practitioners from NMHSs. Interviewees were identified by local collaborators and through snowball methodology, whereby interviewees were asked to identify 2-3 additional stakeholders. We determined that we had effectively sampled all key stakeholders when no new names were identified; this was feasible given the relatively small size of the climate and health sectors in the study area.

A survey instrument was developed for health sector stakeholders. We asked questions regarding basic demographic information, their perceptions of climate variability and arbovirus risk factors, perceptions of the public health sector response to climate variability, current use of climate information, how they prefer to interact with and receive EWS information, the current strengths and weaknesses of their department with respect to the implementation of a arbovirus EWS, and their training needs. In the survey, we defined climate variability as, “short-term changes in climate that occurs over months, to seasons, to years. This variability is the result of natural, large-scale features of the climate, often related to El Niño or La Niña events. Examples include floods, multi-year or seasonal droughts, heat waves, hurricanes or tropical storms.” Questions were informed by a prior large-scale survey of health practitioner perceptions of climate change impacts on health conducted in the United States, called “Are We Ready?” [36–38], as well as studies by Paterson et al. [39] and Gould and Rudolf [35].

Printed surveys were distributed to health sector stakeholders at national vector control, environmental health, and epidemiology offices, as well as those who participated in national workshops on the development of climate services for arboviruses in Barbados and Dominica in April 2017. The workshop in Dominica was organized by the CIMH and the Ministry of Health and Environment (6 health sector participants). The workshop in Barbados was organized by the PAHO and the CIMH (∼21 health sector participants). Survey responses were entered into an online digital database using Qualtrics and responses were tabulated.

An interview instrument was developed for stakeholders from the climate and health sectors. Questions in the interview and survey were similar so that we could triangulate and validate the responses. We also asked which organizations they had partnered with to manage vector borne diseases, which organizations they would like to partner with, how climate and health fit within their current institutional priorities/mandates/competencies, and what strategies would stimulate collaboration between the climate and health sectors. In interviews with program directors, we asked additional questions about available climate and arbovirus/vector data, as well as arbovirus and vector surveillance and control strategies, and climate monitoring and forecasting (see institutional competencies in S1 Text).

Project investigators interviewed stakeholders from the climate and health sectors through in-person meetings or via Skype in April and May 2017. Interviews were audio recorded following permission from interviewees. Recordings were transcribed and coded by project investigators to identify responses by research topic, country/region and sector [53,54].

During the Barbados national workshop, we conducted an exercise where health (n=∼21) and climate sector (n=6) participants were divided into small groups that included representatives from both sectors. Groups were asked to respond to different forecast scenarios (2 week, 3 month, and 1 year forecasts of *Aedes aegypti* larval indices and dengue incidence). They were asked to identify the actions that they would take in response to alerts at each time scale, and they discussed the utility of a vector versus disease forecast. As with interviews, responses were audio recorded, transcribed, and coded.

The survey, interview, and workshop instruments were reviewed and tested by local collaborators, as well as the research team, prior to implementation. Instruments are available in S2 Text and S3 Text.

## Results

We surveyed 32 individuals from the health sector and interviewed 41 individuals from the climate (n=10) and health (n=31) sectors. Respondent demographics are shown in Table 2. Several individuals participated in both interviews and surveys; however, the exact number is unknown since identifiable information was not collected from surveys.

**Table 2.**
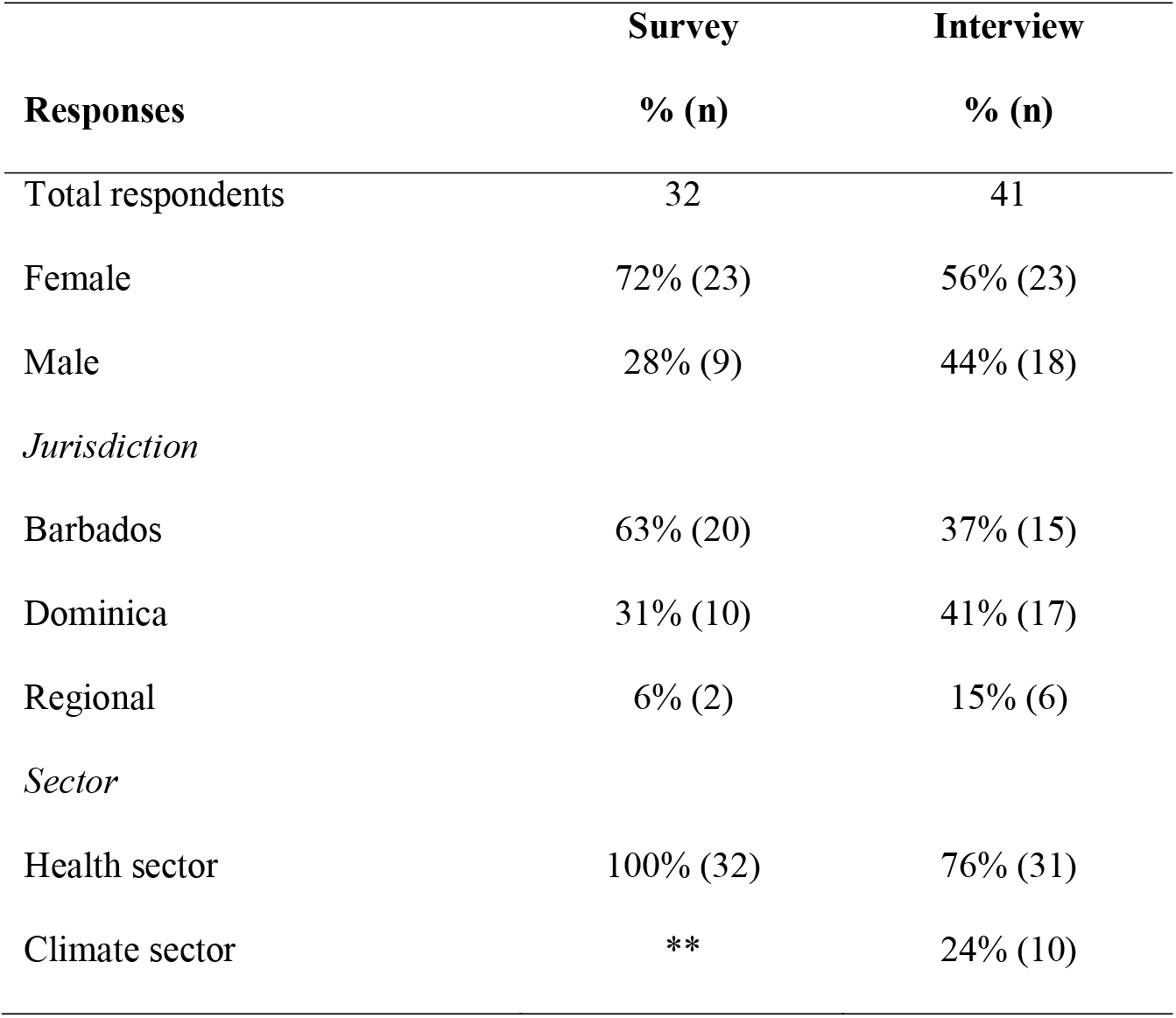
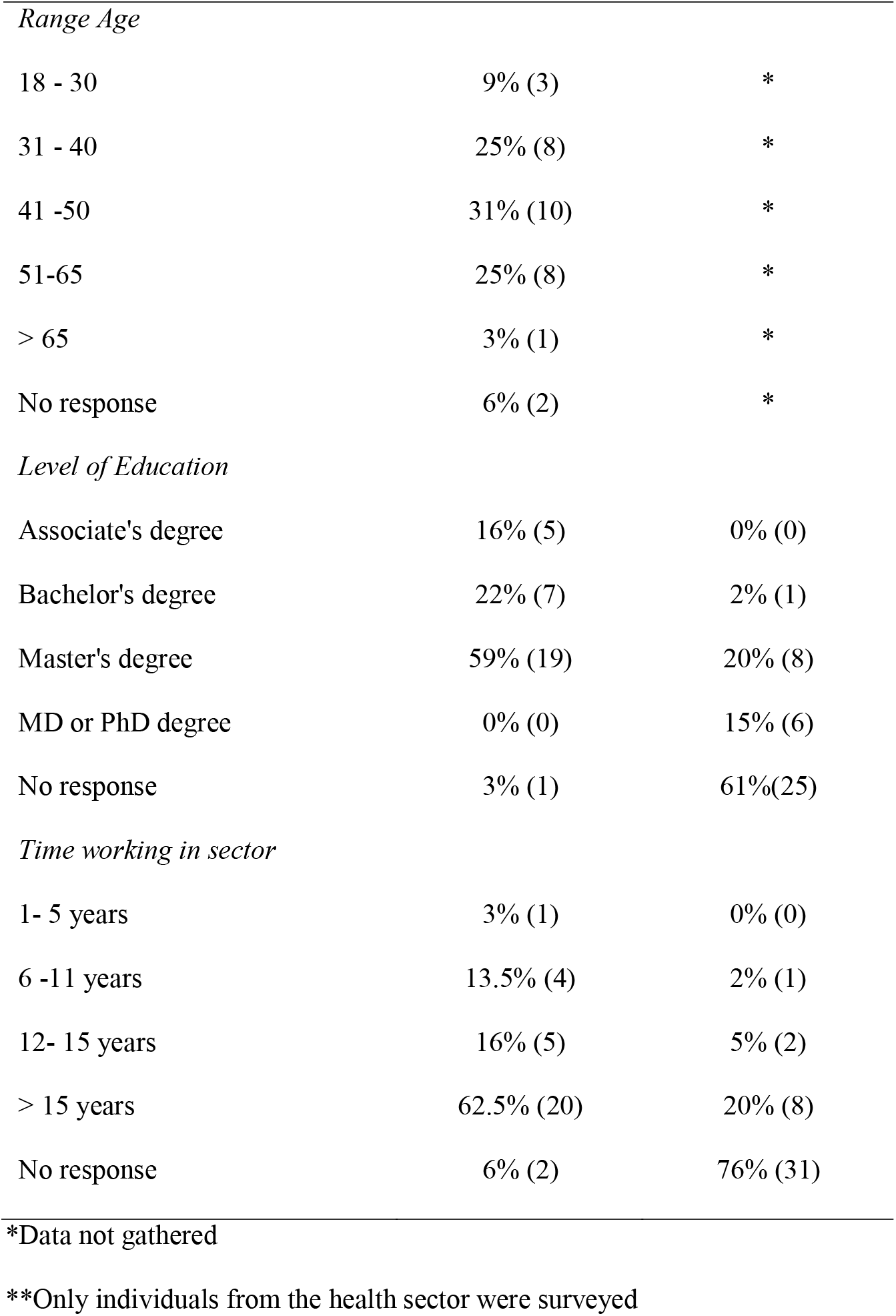
Demographics of survey and interview participants.

|  | Survey | Interview |
| --- | --- | --- |
| Responses | % (n) | % (n) |
| Total respondents | 32 | 41 |
| Female | 72% (23) | 56% (23) |
| Male | 28% (9) | 44% (18) |
| *Jurisdiction* |  |  |
| Barbados | 63% (20) | 37% (15) |
| Dominica | 31% (10) | 41% (17) |
| Regional | 6% (2) | 15% (6) |
| *Sector* |  |  |
| Health sector | 100% (32) | 76% (31) |
| Climate sector | ** | 24% (10) |
| *Range Age* |  |  |
| 18 - 30 | 9% (3) | * |
| 31 - 40 | 25% (8) | * |
| 41 -50 | 31% (10) | * |
| 51-65 | 25% (8) | * |
| > 65 | 3% (1) | * |
| No response | 6% (2) | * |
| *Level of Education* |  |  |
| Associate's degree | 16% (5) | 0% (0) |
| Bachelor's degree | 22% (7) | 2% (1) |
| Master's degree | 59% (19) | 20% (8) |
| MD or PhD degree | 0% (0) | 15% (6) |
| No response | 3% (1) | 61%(25) |
| *Time working in sector* |  |  |
| 1- 5 years | 3% (1) | 0% (0) |
| 6 -11 years | 13.5% (4) | 2% (1) |
| 12- 15 years | 16% (5) | 5% (2) |
| > 15 years | 62.5% (20) | 20% (8) |
| No response | 6% (2) | 76% (31) |

\*Data not gathered
\*\*Only individuals from the health sector were surveyed

### (1) What are the perceptions of climate-health or climate-arbovirus linkages?

#### Perceptions of climate variability and health impacts

In surveys, health practitioners were asked to respond to a series of statements about the effects of climate variability on health in their jurisdiction and their ability to respond to these effects (Likert scale, from strongly disagree to strongly agree, Table 3). Most agreed that their jurisdiction is experiencing an increased risk of diseases transmitted by *Ae. aegypti* due to climate variability and that the risk will increase in the future. Survey respondents were worried about the effects of climate variability on health, and they agreed that this is an urgent problem in their jurisdiction. Although two thirds agreed that there are options or solutions to reduce the effects of climate variability on health, they disagreed that they had sufficient resources and expertise assess the impacts of climate variability on health and to protect residents in their jurisdiction.

**Table 3.**
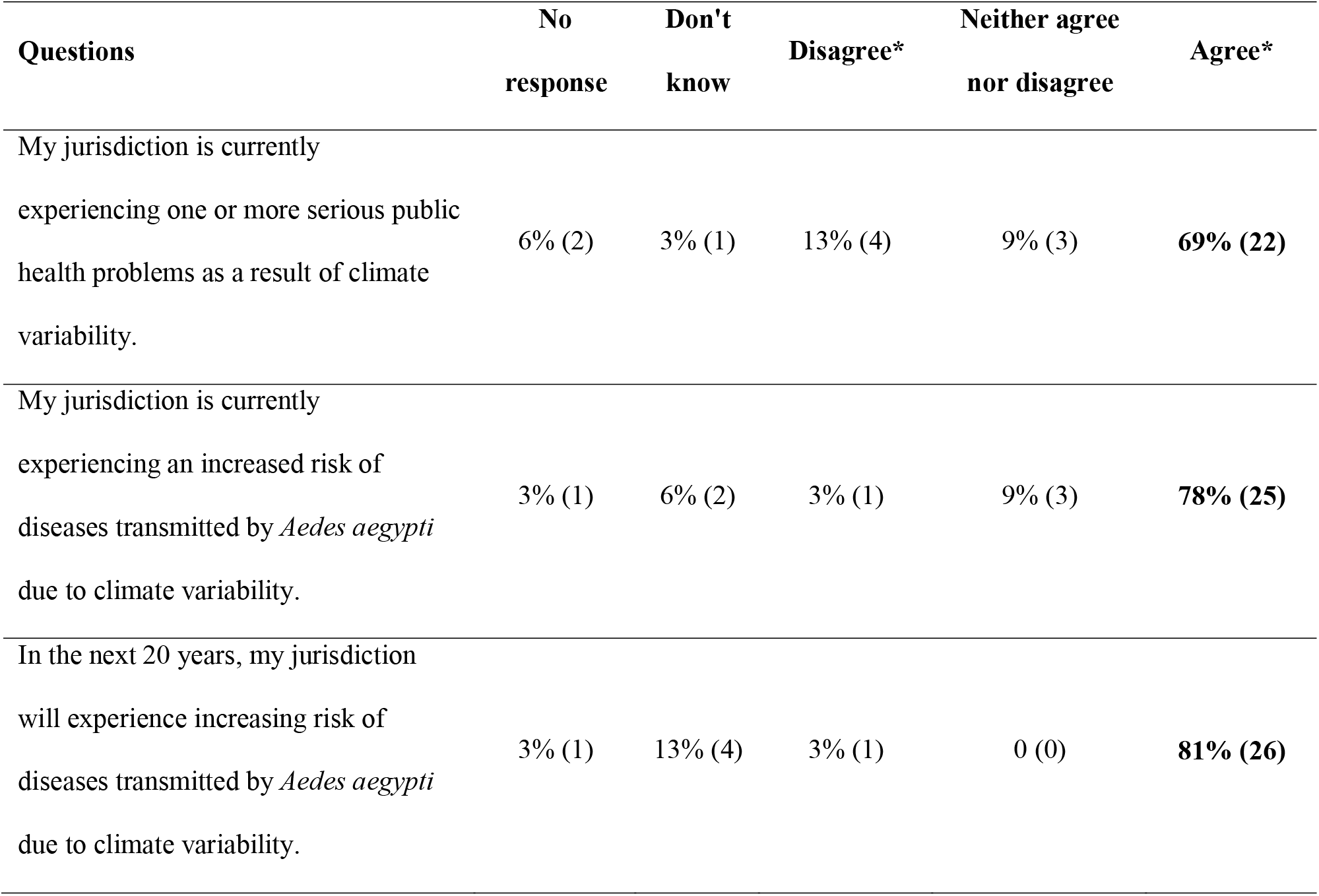
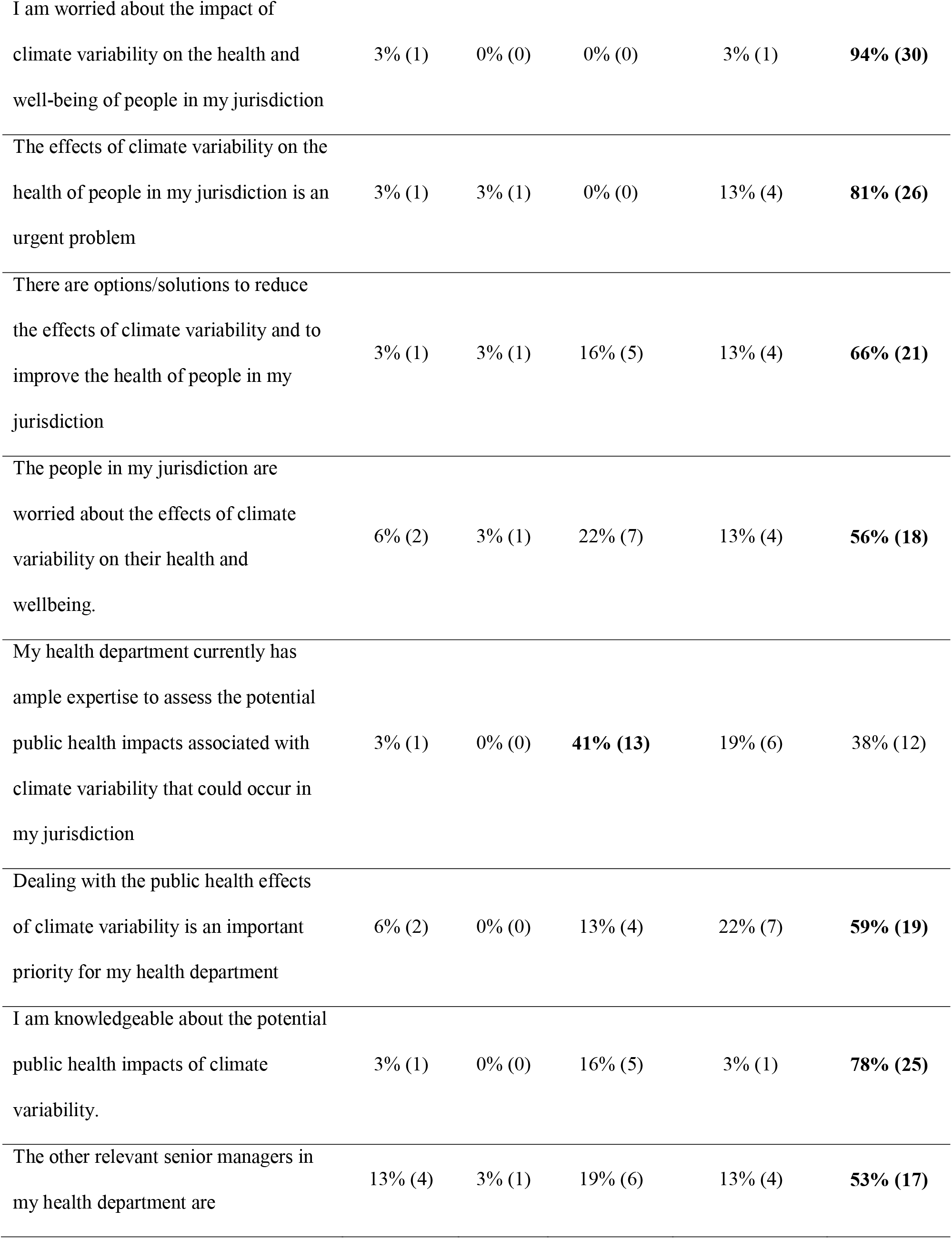
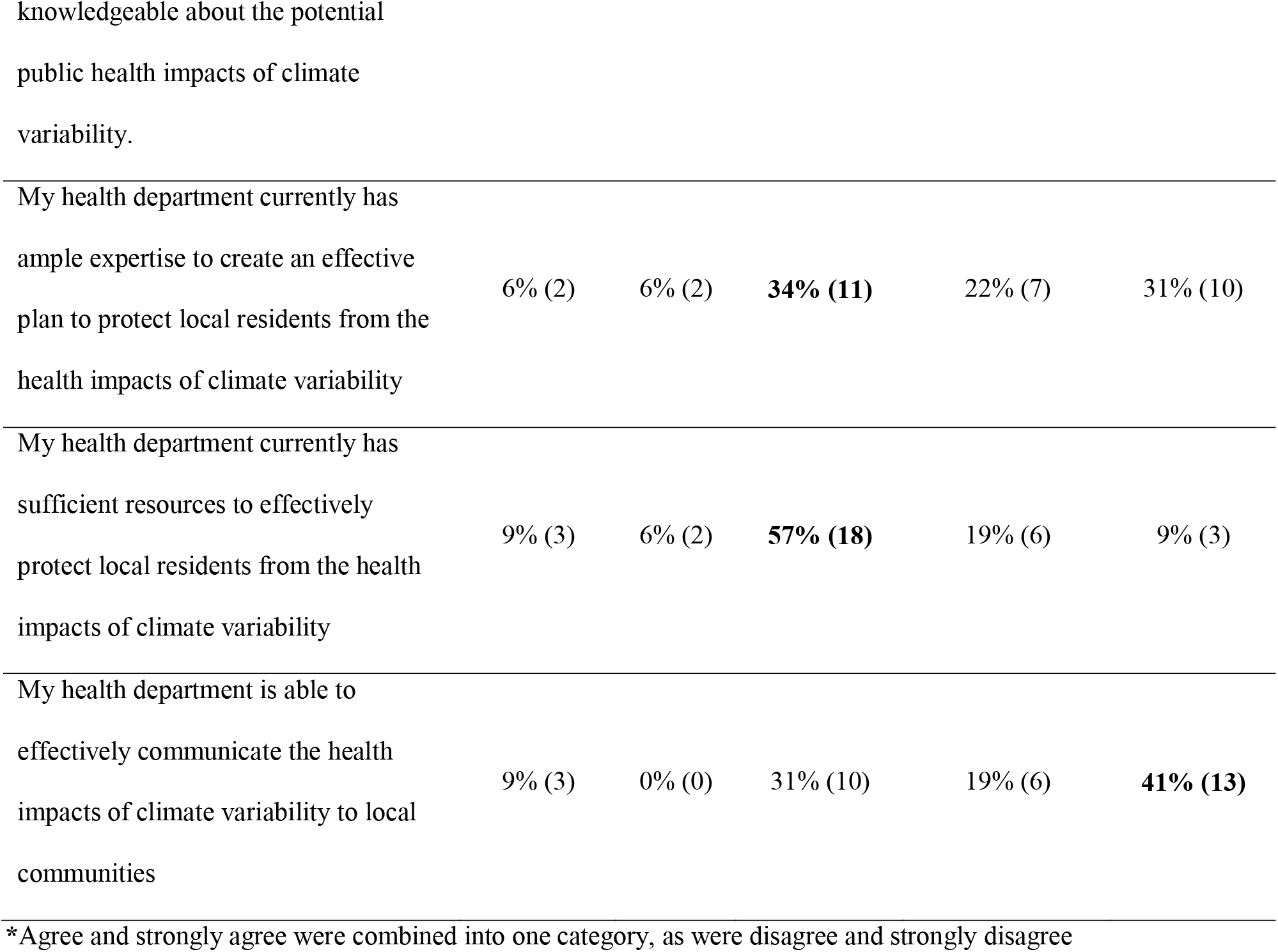
Perceptions of climate variability impacts on health reported by survey respondents. Results shown as % (n). This most common response per question is highlighted in bold. Adapted from [36–38].

#### Climate and non-climate risk factors for Aedes-transmitted diseases

Survey respondents were asked to rank climate and non-climate risk factors for epidemics of diseases transmitted by *Ae. aegypti* (Table 4). Non-climate risk factors were identified as more important overall than climate risk factors. The most important non-climate risk factors, in order of importance, were the introduction of new viruses to susceptible populations, water storage behavior, and insecticide resistance in mosquitoes. The most important climate risk factors, in order of importance, were heavy rainfall and drought conditions. The least important risk factors were warm air temperatures, El Niño or La Niña events, and economic barriers to mosquito control by households.

**Table 4.**
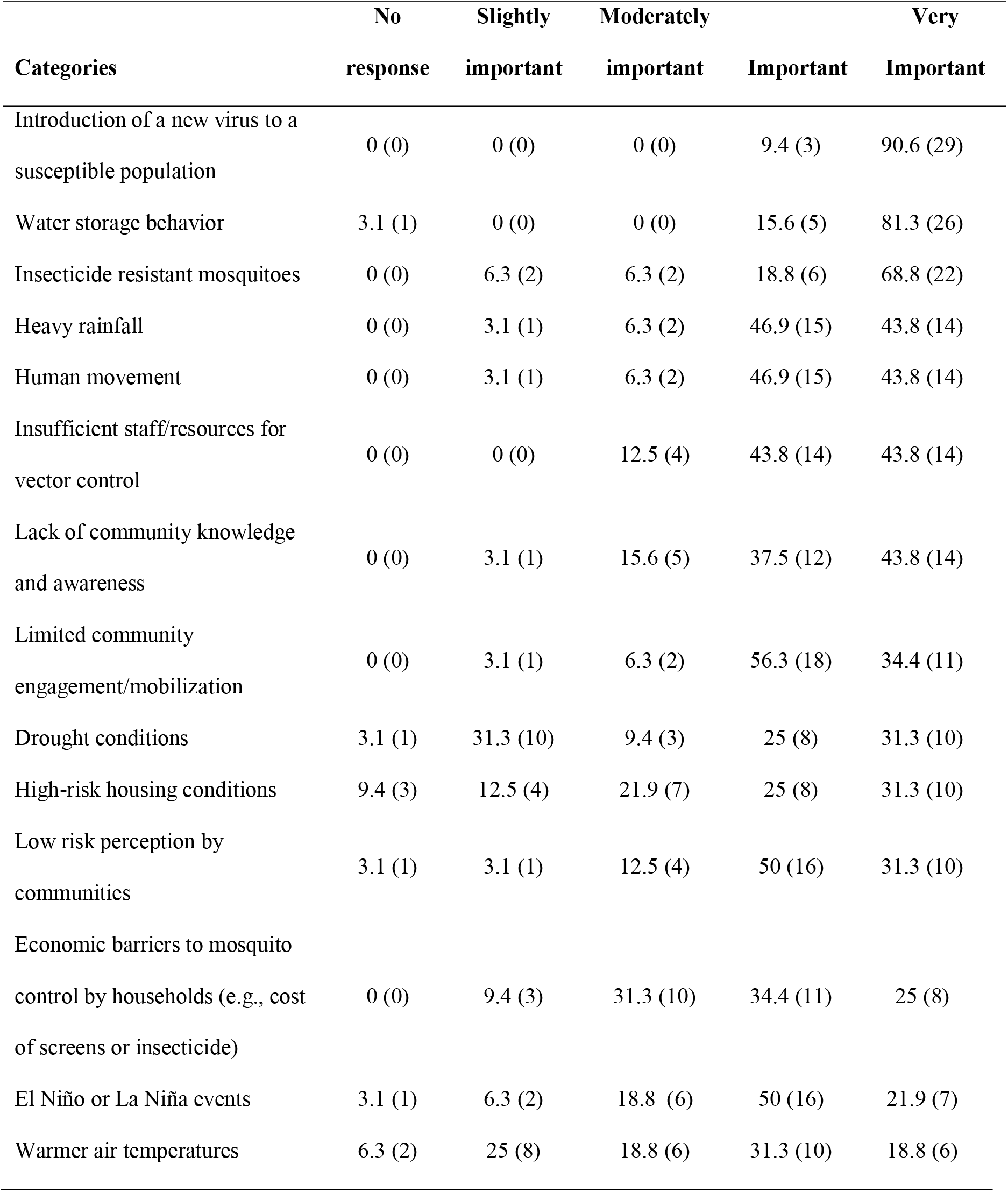
Factors that trigger epidemics of diseases transmitted by Aedes aegypti reported by survey respondents. Results shown as % (n), listed in order of most to least important.

Interviewees were also asked to discuss climate and non-climate risk factors for arbovirus epidemics. They indicated that frequent (re)-introduction of viruses and vectors was associated with human movement between the islands due to trade and tourism. In Dominica, interviewees also identified human movement between rural and urban areas as a risk factor.

Interviewees identified the onset of the hot, rainy/wet season as a risk factor for arbovirus transmission, although they indicated that the linkages between rainfall and dengue fever have become less apparent due to water storage practices. Two interviewees highlighted this contradiction,

> “If the rain falls very heavily, within two weeks expect to have an increase in number of cases. It’s always associated with rainfall” (Health Sector, Barbados). “With these droughts, there doesn’t seem to be, in the last few years, a real dengue season” (Health Stakeholder, Barbados).

In Barbados, interviewees indicated that household water storage was associated with drought conditions and the resulting water scarcity. Another risk factor was the national legislation requiring that all new buildings greater than 1500 square feet have rainwater storage receptacles as a drought adaptation strategy; however, the receptacles have become mosquito larval habitat. Interviewees indicated that the improper management of public utilities and infrastructure (e.g., telephone junction boxes, manhole covers, public wells, drains) had resulted in cryptic mosquito larval habitats that were difficult to find and treat during the wet season.

In Dominica, interviewees commented that water storage had increased following Tropical Storm Erika (2015). The storm damaged the piped water systems and people began storing freshwater in 55-gallon drums around the home. This behavior continued despite repairs to water systems. One interviewee described the effects of Erika,

> “After the Tropical Storm Erika, everything just got a little more vulnerable than it used to be… it was just one downpour of rain that caused all of the destruction” (Health Stakeholder, Dominica)

Interviewees from Dominica noted that *Ae. aegypti* had expanded its range into higher elevation areas, where the mosquito had not been present historically.

#### Other effects of climate on health

Interviewees were asked to identify other ways that climate affected health in their jurisdiction. They identified a wide range of interrelated health effects associated with climate, including increased risk of morbidity due to the interaction of heat stress and diabetes associated with hotter days and nights, leptospirosis (*Leptospira* sp.) associated with flooding, and communicable diseases associated with relocation and crowding of people in shelters following tropical storms. They indicated that malnutrition was associated with droughts that reduced crop yields and warming ocean temperatures that caused fish kills. Respiratory problems (e.g., asthma) were associated with dry weather, dust and air pollution. Factors unique to Barbados included hypertension due to sea level rise and salt-water intrusion in the groundwater supply, reduced hygiene and *Pseudomonas* infections due to water scarcity and storage, skin cancer due to UV exposure, and water-borne diseases (e.g., gastroenteritis, *Salmonella*) associated with flooding. Factors unique to Dominica included loss of lives due to landslides associated with tropical storms, gastroenteritis associated with dry weather, and mental health morbidity in the elderly and other vulnerable populations who are relocated after tropical storms. One informant described the complex web of causality associated with the effects of climate on health,

> “[During droughts] people are not able to go to their farms, they don’t have food and their nutrition suffers. They don’t have income… they cannot get their medications… So its just the rippling effect” (Health Stakeholder, Dominica).

Interviewees stressed the need to strengthen the evidence base linking climate and health in their jurisdictions. They recognized that most of these linkages were anecdotal or hypothetical, since there have been few local studies on climate and health, as summarized by one interviewee,

> “So we have not been able to make a direct link between those diseases and climate variability and change; however, we know that there has been an increase as a result of climate variability… The data to make that linkage… is not really always available” (Health Stakeholder, Dominica).

Interviewees from the Barbados NMHS indicated that they had limited experience with climate research, and that this was an area that they were interested in expanding. National-level health sector interviewees displayed a high level of field experience and local knowledge, but they indicated that they had little knowledge of empirical studies that could inform their decision-making and planning processes. As stated by one interviewee,

> “We want more evidence-based decision-making. We want data… That’s priority #1… to get the evidence.” (Health Stakeholder, Regional)

Interviewees recommended conducting case studies, or demonstration projects, in the region to generate local evidence on climate-health linkages. They suggested focusing these investigations and interventions at the medium-term climate variability timescale (e.g., seasonal variation, year-to-year variation in extreme climate events), rather than the long-term climate change time scale.

### (2) Who are the key actors engaged in climate-arbovirus surveillance and control, and how to strengthen communication and partnerships amongst these actors?

#### Partnerships

Interviewees identified diverse national and international agencies and funders engaged in climate-arbovirus surveillance and control (Figure 2). The key regional institutions were the PAHO, the CARPHA, and the CIMH. The Red Cross was the most frequently mentioned non-governmental organization (NGO). The health sectors engage periodically with their respective NMHS on specific projects; however, there are no formal collaborations. Understanding and mitigating the effects of climate on health are relatively high priorities in the health sector, but climate and health is not yet a mandate (S1 Text). Similarly, the NMHS do not have a mandate to work on climate and health. As a result, it has been difficult to allocate resources (e.g., personnel, funding) to this area. Interviewees indicated that the key partnerships to be strengthened were the private sector (tourism, vector control companies, media), academic institutions, and civil society organizations.

**Figure 2.**
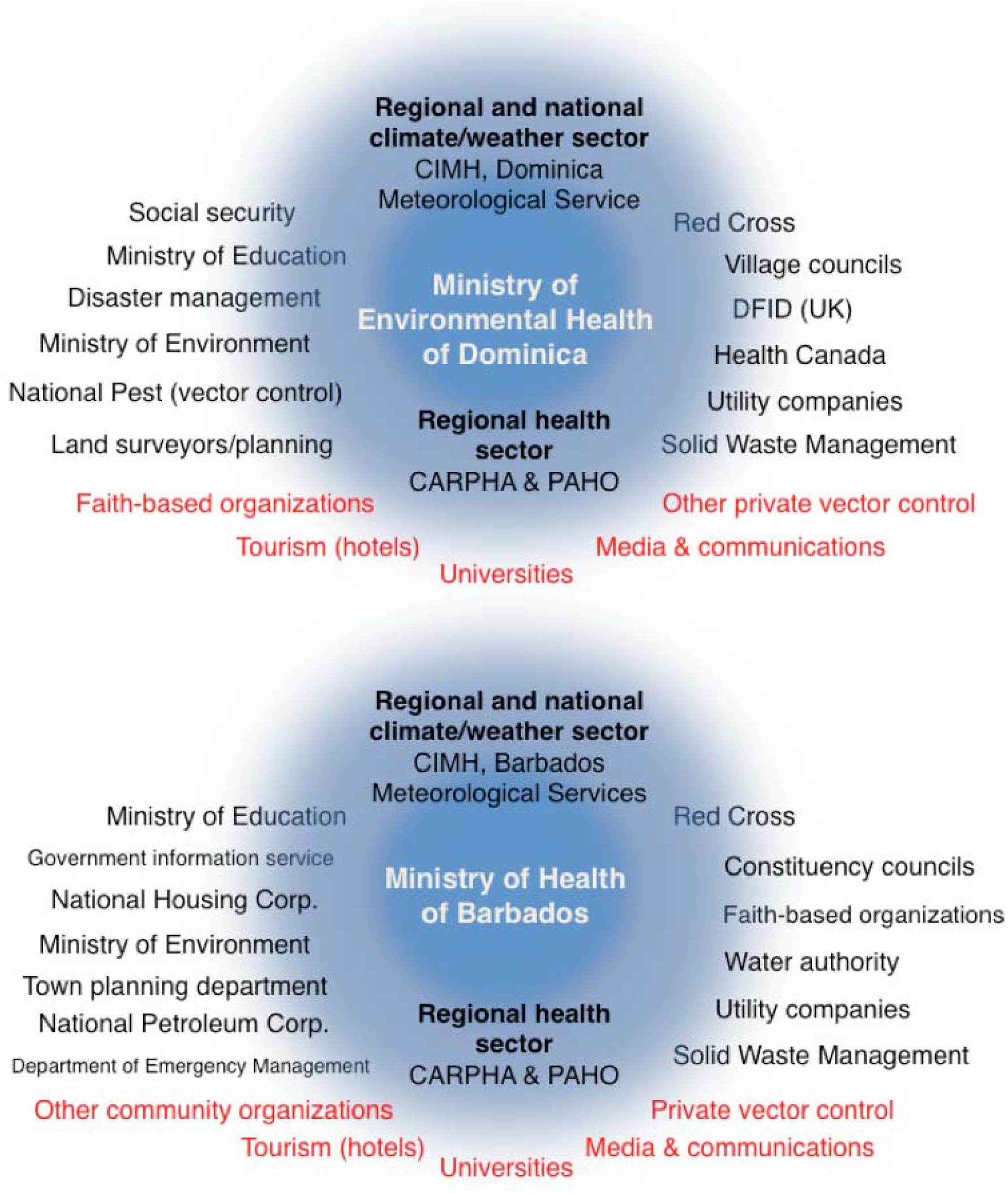
Stakeholder analysis. Organizations (in black) that work with the health sector in Barbados and Dominica on issues related to vector control and climate services for health. Organizations in red were identified as needing a stronger relationship with the health sector.

#### Collaboration strategies

Interviewees identified six strategies to strengthen the communication and partnerships amongst these actors. First, they highlighted the importance of an integrated approach to the development of climate services for health spanning research, operations, a platform for data and knowledge sharing, outreach, awareness raising, education, an in-country response, and mitigation plans and policies. Second, interviewees emphasized the importance of engaging senior leaders from the health sector to raise the profile of climate and health on the health agenda, and to ensure that actions are driven from the top-down. Third, they highlighted the importance of formal collaboration agreements amongst climate, health, and other sectors, similar to the multi-lateral agreements recently signed amongst the CIMH, the CARPHA and other regional Caribbean agencies. Interviewees indicated that collaboration agreements would allow them to co-develop and co-deliver climate services for the health sector.

Interviewees perceived that collaboration agreements signaled a strong commitment from institution directors and an understanding of mutual benefit. Fourth, they suggested that national committees on climate and health be established to specify the work that would be done jointly, the roles of each partner, a timeline for an operational plan, and standard operating procedures (SOPs) with a framework for communication, data sharing, and reporting guidelines. Fifth, interviewees indicated the importance of creating shared spaces for dialogue between the climate and health sectors, such as regional and national climate and health forums. An interviewee from the climate sector stated,

> “Just sitting with people in the sectors makes such a big difference… Understand them, what drives them, what are their needs? Because we might think they need something they don’t… Sometimes it’s about forgetting yourself and putting yourself in the other person’s shoes to really figure out what the need is about. That’s true engagement” (Climate Stakeholder, Regional).

This engagement would help to build functional working relationships and increase the trust among people in both sectors, allowing sectoral stakeholders to learn about the needs and perspectives of the other, what information can be shared, and the resources available to help each other. One interviewee stated,

> “Once we build the trust, then we build the network, then we can see what the willingness to collect, to centralize, to digitize, and to share the data really is” (Climate Stakeholder, Regional).

For example, interviewees suggested that the MoH could partner with their NMHS to ensure that new weather stations are placed in areas that are strategic for the surveillance of arboviruses. Representatives from the NMHS could participate in the regular epidemiological surveillance meetings of the MoH. Last but not least, interviewees suggested that climate services for health be framed as a national development priority, a strategy that would increase buy-in from decision makers and funding from international development agencies. One informant stated,

> “I think people will embrace climate and health… [it is] a real sustainable development goal… Health has always been a critical sector” (Climate Stakeholder, Regional).

### (3) What are the current capabilities of the health and climate sectors to implement a climate-driven arbovirus EWS? What capacities need to be strengthened?

Health sector survey respondents were asked to identify the strengths and weaknesses of their institution with respect to the implementation of an arbovirus EWS (Figure 3). The top strengths were effective public health messaging to communities, effective health surveillance infrastructure, knowledge of the effects of climate on vector borne diseases, strong coordination with other institutions, and community mobilization. The top weaknesses, or areas to be strengthened, were the availability of financial resources and expertise in geographic information systems (GIS), statistics, modeling, and computer programming (S4 Table for software currently used in health departments and S5 Table for preferred training activities).

**Figure 3.**
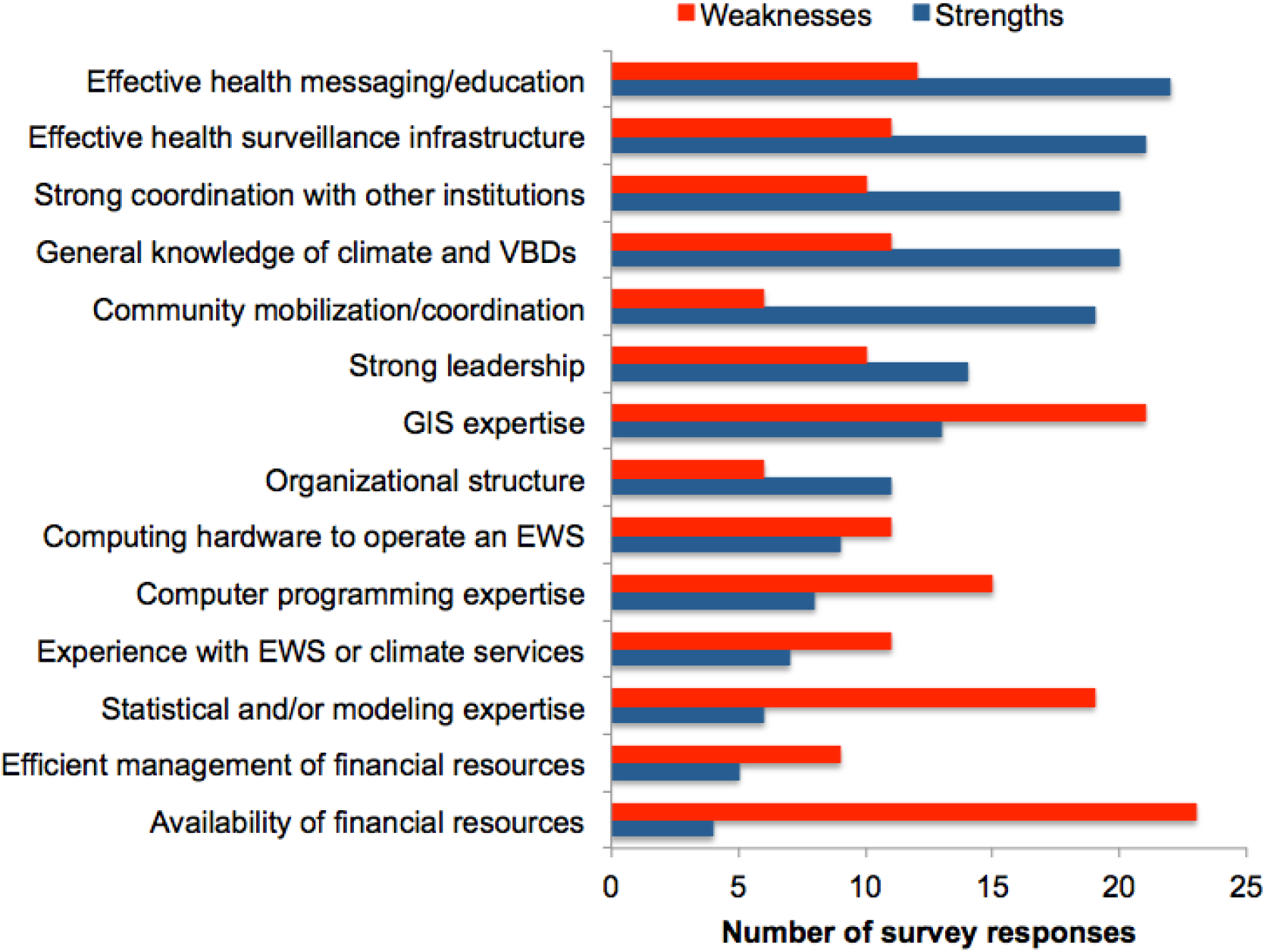
Perceptions of the strengths and weaknesses of the health sector with respect to the capacity to implement an EWS for *Aedes aegypti* transmitted diseases. EWS = early warning system, GIS = geographic information system, VBDs = vector borne diseases

Interviewees highlighted the need to train, nurture, and retain a cohort of practitioners with expertise in climate and health. Health sector interviewees emphasized the need to increase their skills in modeling and data analysis through technical workshops on how to use climate information, data, models and other tools to predict epidemics. They identified the need for training on climate and health linkages, greater understanding of climate services for health, how to use climate services for health during emergencies/disasters, and how to communicate the effects of climate on health to local communities. They suggested that training activities be practical and interactive, such as workshops where multisectoral teams respond to simulations of epidemic warnings.

Health sector interviewees also expressed an urgent need for training on geographic information systems (GIS). They found that GIS was a highly effective tool that allowed them to use field data to make informed decisions, and to communicate risk information back to the public.

> “It is so much easier, better, to use maps when you are doing presentations. Especially if you are doing something with the public where you can actually show them their community and say, ‘There you have breeding sites. There is where you have the problem.’ And they can actually see it. You can actually show it to them.” (Health Stakeholder, Dominica).

In Dominica, interviewees highlighted the need for vector control specialists in the MoH, since their environmental health officers are responsible for a broad portfolio of activities and are trained as generalists. Interviewees from both countries highlighted the need for better data collection and storage practices in the health sector in order to create high-quality, long-term datasets.

NMHS (national climate sector) interviewees indicated that they had limited capacity to implement climate services for health. Interviewees from Barbados and Dominica indicated the need for additional personnel, as well as a better understanding of health sector end-user needs Interviewees from the Dominica NMHS indicated that they had difficulty reaching the meteorological stations to download data due to the complex topography of the country. They identified the need for basic resources to increase their monitoring and forecasting capacities, including a staff meteorologist, adequate transportation to and from meteorological stations, financial resources, instrumentation, and improved security to prevent vandalism the meteorological stations. They stated,

> “We use our personal vehicles, but some of the areas are a bit challenging, and we are two females, so sometimes …depending on where we are going, we need somebody else to go with us, for security” (Climate Stakeholder, Dominica).

Health sector interviewees from Dominica suggested that their technicians could be trained to download data from meteorological stations to support the local NMHS.

### (4) What climate/weather data are currently used by the health sector for arbovirus control, what is the added value, and how can climate/weather data be effectively mainstreamed for arbovirus control operations?

#### Use of climate information

Health sector survey respondents were asked about their current use of climate information (S6 Table). Two thirds of respondents indicated they had received general information on the effects of climate on vector borne diseases, and half of the respondents confirmed that climate information was used for some level of planning for disease and vector control interventions. They were unsure as to whether an EWS for arboviruses existed in their jurisdiction, but most indicated that climate information was not part of existing epidemiological warning systems.

Interviewees confirmed that the arbovirus alert systems in Dominica and Barbados were based solely on epidemiological surveillance. The health sector and PAHO issue an alert when the number of reported cases surpasses a pre-determined threshold established by the historical average for the same week or month (see Lowe et al. [11] for details). Interviewees indicated that the current system did not provide sufficient lead-time to effectively reduce the threat of an epidemic.

Interviewees described the current use of climate information for arbovirus control. The health sector considers wet/dry seasons and extreme climate events when planning vector control programs, for example, by increasing larviciding efforts at the onset of the wet season or increasing community campaigns on safe water-storage during droughts. Occasionally the health sector requests climate/weather information from their NMHS and the data are generally shared as Excel files. However, they do not formally incorporate climate information, such as seasonal climate forecasts, into their planning process. Overall, climate information was reported to play a minor role in decision-making, which was instead driven by policies, regulations, and specific competencies of the organizations.

#### Forecast scenarios

At the national workshop in Barbados, health and climate stakeholders were asked to identify the interventions they would implement if they were provided with short (2 week), medium (3 month) and long-term (1 year) forecasts of vector abundance and dengue incidence (S3 Text). They unanimously stated that disease incidence forecasts would be more effective than vector forecasts in garnering the political attention necessary to mobilize resources to implement preventative interventions. With a short-term forecast, the health sector would increase education, community mobilization, and larval source reduction, especially in known hotspots. With a medium-term forecast, the health sector would be better able to plan with stakeholders, mobilize the field team, look at trends, and create bulletins for community mobilization. With a long-term forecast the health sector could better lobby with health sector leadership and the Minister of Finance for the needed financial support, allowing for more effective budgeting. They would be able to monitor and evaluate interventions and conduct a needs assessment to inform planning. They would also be able to procure diagnostic reagents and supplies for the national reference laboratory, a process that can take up to 6 months. Although the workshop participants identified meaningful interventions at each timescale, they preferred the medium-term (3-month) forecast as indicated in the following,

> “A year can feel like a long time away. With 3 months, there will be a sense of urgency and you can do meaningful activities, although there might not be new resources” (Health Stakeholder, Barbados).

#### Added value of climate services

Health sector interviewees highlighted the ways in which climate services would improve their planning for arbovirus interventions. By integrating climate and/or disease forecasts into their seasonal and annual planning processes, they felt they could be proactive and more effective at preventing outbreaks, as described by this interviewee,

> “We know we have Aedes, we know we have the threat, but its only when outbreaks happen, we start scrambling around to do things. So I think if we can put mechanisms in place, long in advance, then we can have more success in dealing with outbreaks. Or we can even prevent outbreaks” (Health Stakeholder, Dominica)

Interviewees indicated that climate services have to provide reliable information early enough such that the health sector can target control efforts in high-risk areas during certain times of the year. This would result in a more efficient use of limited financial and human resources, as described by this interviewee,

> “When you know that there is an impending threat, you would come up with specific activities that you would conduct. It doesn’t necessarily mean that those activities would be at a higher cost, but you can be more specific… It will be easier for us to respond to an impending threat, instead of running around” (Health Stakeholder, Dominica).

Interviewees indicated that forecasts of disease risk could be used to inform hospitals about staffing needs, stocking of medicines and laboratory diagnostic reagents, and the development of targeted educational materials for the public. They suggested that warnings be communicated to the public through social media and other outlets to motivate community mobilization for preventative practices. Interviewees indicated that they would feel more motivated and inspired in their day-to-day work if they could see how the data that they collect was being used to inform decision-making.

#### Mainstreaming climate services

Health sector survey respondents were asked how they would prefer to receive information from an arbovirus EWS. The top responses were: a climate and health bulletin (91%), an interactive GIS platform (66%) and internal meetings within their departments (59%) (S7 Table).

Interviewees were also asked to identify climate services that would improve their day-to-day work related to arboviruses. Health sector interviewees confirmed that they were interested in utilizing the Caribbean Health Climatic Bulletin launched in 2017 by the CIMH, the CARPHA and the PAHO. The bulletin qualitatively summarizes potential health impacts for a 3-month period based on seasonal climate forecasts (https://rcc.cimh.edu.bb/health-bulletin-archive/). Health sector interviewees also reiterated that a GIS platform would allow them to integrate and analyze real-time information on disease epidemiology, entomology, and climate. They could use this platform to produce risk maps showing the spatial distribution of mosquito vectors and disease risk in relation to rainfall, temperature, and other climate information. They suggested that these forecasts be converted into spatiotemporal alerts using a color-coding scheme. Other ideas for climate services included the use of wind speed and wind direction forecasts to inform insecticide fogging operations. One informant summarized the health sector needs in the following,

> “We need it [climate services] packaged in such a way that the health professional would understand. Pick it up, and look at it, and understand it” (Health Stakeholder, Dominica).

> “Decision makers at the policy level are not healthcare providers. They are administrators, they are politicians, and we need to help them. We need to feed them [decision makers] with the kind of information they can understand, and [so] they can feel comfortable making decisions” (Climate and health workshop participant, Barbados).

## Discussion

Small island developing states (SIDS) in the Caribbean region are amongst the most vulnerable countries in the world to climate change [1,2]. The effects of a changing climate include increased frequency and severity of droughts and increased frequency of tropical storms and hurricanes. Extreme climate events affect directly or indirectly most dimensions of people’s well-being, including mental and physical health, food, housing, freshwater, and livelihoods. Huang et al [55] state that public health sector adaptation to climate change should consist of both adaptive-capacity building and implementation of adaptation actions. A climate-driven arbovirus EWS is a key adaptation action. However, our study confirms that this will only be possible if current capacities are increased in the Caribbean region. Prior reports have noted the embryonic status of the application of climate science in the Caribbean health sector [24] and an assessment of on the capacity of the network of Caribbean NMHSs to deliver climate services that found that only one NMHS offered specialized climate information services for the health sector [56].

Global research on climate services has identified characteristics or conditions to develop a usable science that can be mainstreamed for public health operations. The first phase is to establish an enabling environment for partnership with different stakeholders. This is done by identifying the common priorities, needs for research, and by building necessary capacities for understanding among climate and public health stakeholders and researchers [57,58]. In this study we identified some of the challenges involved in initiating a successful process of joint collaboration between the climate and public health sectors. This partnership is critical to ensure commitment and ownership by different stakeholders and end-users as climate services are developed.

In this study, climate services for health, specifically for *Ae. aegypti*-transmitted arboviral diseases, were considered because high policy level organizations (PAHO/WHO, CIMH, CARPHA) are encouraging the application of the GFCS. The WHO recently developed a guide for the operational development of climate resilience water [59] and health systems [60]. This initiative acts in synergy with the GFCS and describes how existing systems can include climate information in their operations to reduce the impacts of climate change on human health. We found that the sustainability of these initiatives will require the political will to establish climate services for health as a mandate in the NMHS and public health sectors, allowing them to work in interdisciplinary and intersectoral teams. The benefits of these decisions could extend to other government agencies and the private sector, such as the Ministry of Environment or Ministry of Tourism, as well as regional universities, research centers, and sustainable development agencies. A clear opportunity was the “Third Global Conference on Health and Climate: Special Focus on SIDS,” which was held in Grenada in October 2018. The meeting convened Caribbean Ministers of Health, Ministers of Environment, representatives from UN agencies and other key stakeholders to develop an Action Plan on Health and Climate Change for the Caribbean [61].

We found that the national climate sectors (NMHSs), in particular, identified capacity challenges when asked about engaging in work on climate and health. We also found that the regional climate stakeholders were more experienced with climate services for health than local NMHS stakeholders. Prior assessments of the NMHSs in the Caribbean also noted that there is limited technical capacity of local level staff, especially on smaller islands like Barbados and Dominica [56]. In recent research, representatives from the NMHSs recommended transitioning from a designation of NMHS to become “National Climate Services Centers (NCSC),” which would facilitate the development of climate services at the national level [56].

Despite major advances in climate science and climate-health research globally, our results confirm that climate information is neither routinely applied nor used in planning interventions in Barbados or Dominica. To date, there has been limited success in developing operational climate services for dengue, although several studies have demonstrated the potential [17,62–68]. One of the most promising examples of a climate-driven dengue forecast model framework was recently described by Lowe et al. for Barbados [11]. Distributed lag nonlinear models [69] were coupled with a Bayesian hieratical mixed model [66] to quantify the nonlinear and delayed impacts of climate factors, such as drought and extreme rainfall, on dengue risk in Barbados from 1999 to 2016. The study found that drought periods followed by a combination of warm and wet weather several months later could provide optimum conditions for imminent dengue outbreaks. The developed model successfully predicted a high probability of dengue outbreaks versus non-outbreaks in most years, with improved performance during El Niño years. However, model performance in 2015-2016 was compromised by the lack of data on the emergence of chikungunya and Zika in the region in the prior years. Seasonal climate forecasts routinely produced by the Caribbean Institute for Meteorology and Hydrology could be incorporated in the model framework as an early warning tool. This could help the health sector to plan interventions that mitigate the impact of mosquito-borne disease epidemics in the region up to three months in advance.

The results of this study highlight the need for local research on climate-health linkages, particularly at the climate variability time scale. In some cases, policy makers and practitioners may be better able to plan interventions and to allocate resources at this intermediate season-to-season timescale, as compared to the long-term climate change timescale. Forecasts of extreme climate events can inform Disaster Risk Reduction (DRR) interventions as well as health interventions for the most vulnerable groups. Local climate-health research would engender collaborative scientific articles with co-authors from the climate and health sectors, thus facilitating data sharing, building trust, and fomenting a culture of research on climate and health. Development of data sharing protocols between the climate and health sectors is a priority given the sensitivity of sharing health information.

Health sector stakeholders demonstrated concern, awareness and a high-level understanding of the impacts of climate variability on arboviruses and health in general. Prior research in the Caribbean [42] found that there was limited knowledge about climate and health linkages amongst nurses and doctors in private and public sectors. However, this study focused on health sector practitioners and decision makers engaged in environmental health and vector borne diseases, which may account for their greater awareness and concern. More recent studies in the Caribbean confirm a relatively high level of awareness and concern amongst health-care providers [41], similar to studies in the U.S. [36,38]. Several capacity building initiatives undertaken by regional institutions such as CARPHA and PAHO have likely contributed to higher levels of climate-health awareness over time. However, our findings suggest that the public health sector does not feel ready to develop and implement an EWS or other adaptation measures due to limited institutional capacity (resources and expertise), as found in prior studies in the U.S. and Canada [35–39,55].

With respect to expertise, we identified a demand for basic training to increase technical knowledge on climate and health linkages. There is the opportunity to create capacity building programs focused on climate and health, which would support the creation of a cohort of practitioners, decision makers, and researchers who are specialized in this area, thus providing long-term sustainability for a program on climate and health. Interviewees proposed that joint training activities across the climate and health sectors would increase knowledge while strengthening the intersectoral partnership. An interdisciplinary approach is needed for successful implementation of climate services for health.

Our findings suggest that technical skills are most needed in the health sector, including GIS, statistics, modeling and computer programming. The health sector currently uses basic tools for disease mapping. It is important to assess local user–needs in order to develop tailored visualizations that are useful and relevant. This is an opportunity for co-production with health and other sectors such as urban planning, disaster risk management, city utilities and services. Developing targeted training in GIS that is driven by user-needs will help in visualization and data analysis at the local level. Additionally, there is the opportunity to develop a fuller range of user-friendly tools/instruments that can be applied by the health sector without specialized expertise in their routine data and reporting activities. The operational co-production of tools and products, such as the quarterly Caribbean Health Climatic Bulletin by the CARPHA, the PAHO and the CIMH, is a noteworthy first step. The bulletin includes qualitative expert statements on probable health risks associated with seasonal climate forecasts (3 months ahead). However, there is significant scope for the development of the next generation of climate services that focus on quantitative probabilistic forecasts of disease risk [24]. Overall, the appropriate involvement of stakeholders is a key element to identify users’ needs, to develop users’ capacities and to exploit existing capabilities.

### National-level opportunities

With respect to financial capacity, the Caribbean climate and health sectors are beginning to work together to attract the resources needed to increase local capacities to develop climate services for the health sector. Stakeholders recommended framing climate services for health as a national development priority, thereby attracting funding from international development agencies. A high-level policy goal may enhance the partnership between climate and health government institutions.

For example, the Sustainable Development Goals (SDG), and the Paris Climate Agreement are international policies that have common priorities and objectives: good health and wellbeing (Objective 3), climate action (Objective13), and partnership for the goals (Objective 17) [70,71]. The Paris Agreement recognizes the need to strengthen the global response to the threat of climate change and to significantly reduce the risks of climate change (Article 2.1), including the risk to human health [72]. Thus it is critical to develop national policy interactions within the SDGs, to avoid policymakers and public health planners operating in silos [55,73].

The National Development Plan, National Adaptation Plan for Climate Change, and National Disaster Management Plans are policy documents being developed by most countries in the Caribbean. The PAHO has also led recent efforts to develop Health National Adaptation Plans focused on climate resilient health systems for Caribbean SIDS [73]. These documents provide a policy mechanism to establish lines of intersectoral and interdisciplinary work, which can include the development of climate services for health. Climate change adaptation/mitigation measures are part of the National Determined Contributions (NDCs) that nations develop under the Paris Agreement. Many of those measures can generate co-benefits or added value for the health sector; co-benefits are the additional benefits that result when nations act to control climate change [35]. For example, adaptation efforts aimed at improving water management could also reduce the burden of water-borne or vector-borne diseases [59]. However, specific measures need to be identified, monitored and evaluated to be included into the country National Determined Contributions (NDCs) to be reported to the United Nations Framework Convention on Climate Change (UNFCCC). This approach also offer the possibility to access funding from the Green Climate Fund (GCF) in priority sectors such as health, food, water security, and livelihoods of people and communities. For example, the GCF recently approved (March 2018) and co-funded a 5 year project, “Water Sector Resilience Nexus for Sustainability in Barbados” [74].

Beyond the climate and health sectors, we identified a complex web of institutional actors who can engage strategically in the development of climate services for health including: a) Water agencies and their climate change and SDG goals related to water supply, and water quality (drinking and wastewater) because of their potential links to vector- and water-borne diseases, b) disaster risk management agencies that deal with hydroclimatic risks that impact vulnerable populations and key infrastructure, c) tourism, to protect visitor health, as well as considering human mobility a critical factor for disease transmission, d) private sector vector control companies, e) community based organizations, and f) academic partners, such as the University of West Indies. Identifying priorities and gaps in specific information would strengthen the partnership amongst the sectors, making more effective the development of climate services for human health beyond the ministries and offices of public health [75,76].

### Regional-level opportunities

In its role as the Regional Climate Centre (RCC), the CIMH leads the implementation of the GFCS in the Caribbean. In its thrust to develop sector-specific climate information, the CIMH has pursued an interdisciplinary team approach that leverages the synergies offered by lead technical institutions who are intimately familiar with their national, regional and sectoral contexts, and can consistently invest in the co-production of user-driven climate early warning information [56,77]. The Consortium of Regional Sectoral Early Warning Information Systems across Climate Timescales (EWISACTs) Coordination Partners is an inter-institutional alliance for climate resilience that in its form and function reflects good practice that prioritizes cross-sectoral, interagency models of climate service delivery over those that follow a silo-ed ‘build it and they would come’ approach [24]. As of 2015, the CIMH has actively worked on an emerging, multi-pronged health-climate portfolio in collaboration with national and regional partners such as Ministries of Health, NMHSs, the CARPHA, the PAHO, and other international, interdisciplinary research partners [24]. New and emerging research is being conducted to investigate the linkages between climate and vector borne diseases, heat and health, as well as, Saharan dust and health [24]. Work is also being done in climate and agriculture, which supports health and nutrition.

The Caribbean Community (CARICOM), a group of 20 Caribbean countries, has mandated the Caribbean Community Climate Change Centre to mainstream climate change adaptation strategies into the sustainable development agendas (UNESCO, 2017). For the Caribbean Region, regional perspectives and considerations are relevant for all the countries [78,79]. Successful projects and tools developed in pilot projects, as done in Barbados [11], can be replicated in similar setting in other countries. A demonstration of the benefits of climate services for arbovirus interventions in one country can be used as a model for other productive sectors (tourism, water supply, disaster risk management), and other countries in the region. Regional institutions should work in cooperation to build technical capacities and resilient communities across the region. This is already happening through the Sectoral EWISACTS portfolio and the work of its multi-institutional Consortium, is increasing expertise and awareness of users and providers. CIMH plans to strengthen their RCC platform for engaging stakeholders to share lessons and promote awareness of climate services based on user-needs for all sectors.

### Limitations

When comparing the results of this study to prior studies on health sector perceptions of climate, one key difference is that our study focused on people working with arboviruses, environmental health, and climate, whereas other studies focused on health-care providers or public health professionals in general. However, given the relatively small size of the health sector in Barbados and Dominica, we interacted with most senior leadership in interviews and national consultations, in particular those involved with overall management of the public health sector, epidemiological programs, environment, climate change and health. Although we did consider a regional perspective, the results of this study may not be generalizable to all of the Caribbean. Country-level studies should be conducted to capture the nuances of local governance structures, disease epidemiology, and climate.

Our results were skewed towards the health sector perspective rather than the climate sector, given that more health sector stakeholders were interviewed, and only health sector stakeholders were surveyed. In part, this reflected that there were many more people working in the national health sectors than in the national climate sectors. On the climate side, our results were skewed towards the regional perspective, given that regional stakeholders had more experience with climate services for health.

### Conclusion

The results of this study provide recommendations to enhance an interdisciplinary dialogue and partnership within an active community of practitioners, decision makers, and scientists [31,80]. This study contributes to a broader effort to work collaboratively with regional and national health and climate stakeholders in the Caribbean to develop decision support models to predict arbovirus risk and to design effective warning and intervention strategies [24].

One of the key conclusions of this assessment is the need to strengthen the provider-user interface, as currently there is only limited consideration of the products needed by health sector users. Climate services for health can only become operational with the will and support of the climate and health sector institutions. At the same time, it is necessary to create appropriate ‘communities of practices’ and to emphasize the co-design of climate services products [81]. Final recommendations include:

1. To continue to assess local stakeholder needs and perspectives to support the development of climate services for health.
2. To establish a Memorandum of Understanding (MoU) or Letter of Intent (LOI) between the climate and health sectors, particularly at the national level, with a focus on the development of climate services [77].
3. Strengthening the capacity of NMHS through their designation as National Climate Services Centers (NCSC) [56], allowing them to build capacity around the basic and operative aspects of climate services and to collaborate with health sector partners to promote climate services for health.
4. National Adaptation Plans for Climate Change, including recent regional efforts to create Health National Adaptation Plans, may be an opportunity to include a policy or mandate for climate in the health sector, and may be an opportunity to strengthen climate services, applying long-term scenarios for planning in health and other sectors.
5. To strengthen health sector engagement in the region through annual forums focused on climate services and capacity building tailored to the health sector. This could build on existing regional climate meetings like the bi-annual Caribbean Climate Outlook Forum convened by the CIMH.
6. To implement the WHO operational framework for building climate resilient health systems [60], and to implement other mechanisms to integrate climate data, information and knowledge with multiple health and non-health data sources to support decision making.
7. To improve technical GIS and modeling capabilities, and to develop locally relevant tailored tools for non-experts, which can be used to inform decisions and decision-making processes.

## Acknowledgements

This study was solicited by the Caribbean Institute for Meteorology and Hydrology (CIMH) through the United States Agency for International Development’s (USAID, Grant ID: AID-538-10-14-00001) Programme for Building Regional Climate Capacity in the Caribbean (BRCCC Programme: rcc.cimh.edu.bb/brccc) with funding made possible by the generous support of the American people. The authors thank Dr. Shelly-Ann Cox for her data collection and research support. RL was supported by a Royal Society Dorothy Hodgkin Fellowship. The funders had no role in study design, data collection and analysis, decision to publish, or preparation of the manuscript.

## Competing interests

The authors have declared that no competing interests exist.

## Supporting information legends

**S1 Text. Climate and Health Sector Mandates and Competencies.** This document describes the mandates and competencies of regional (Caribbean) and national (Barbados and Dominica) climate and health sectors with respect to arbovirus and vector surveillance and control, and climate monitoring and forecasting. Information was gathered through face-to-face interviews with key stakeholders.

**S2 Text. Interview and survey instruments**. This document contains (1) questions about climate information for arboviral control, used in interviews with climate and health decision makers, managers and expert practitioners, (2) interview questions regarding climate and health data, (3) survey for health sector decision makers, managers, and expert practitioners.

**S3 Text. Forecast scenarios discussed in the Barbados stakeholder workshop.** This activity was conducted at a national consultation at the PAHO in Bridgetown, Barbados, in April 2017, with 27 representatives from the national Ministry of Health (MoH) of Barbados, the Barbados Meteorological Services, the CIMH, and the PAHO. Participants were divided into small groups that included representatives from climate and health sectors. Groups were asked to respond to different forecast scenarios (2 week, 3 month, and 1 year forecasts of *Aedes aegypti* larval indices and dengue incidence). They were asked to identify the actions that they would take in response to alerts at each time scale, and they discussed the utility of a vector versus disease forecast. Results were identified by coding the transcripts of audio recordings.

**S4 Table. Types of software currently used in health departments.** Results from surveys are shown as % (n).

**S5 Table. Preferred training activities identified by health sector survey respondents.** Results from surveys are shown as % (n).

**S6 Table. Current use of climate information and early warning systems reported by survey respondents.** Results shown as % (n).

**S7 Table. Preferred way of receiving information from an early warning system that predicts arbovirus epidemics.** Results shown as % (n).

