## Supplement 1 Text for "Co-developing climate services for public health: stakeholder needs and perceptions for the prevention and control of *Aedes*-transmitted diseases in the Caribbean"

**S1 Text. Climate and Health Sector Mandates and Competencies.** This document describes the mandates and competencies of regional (Caribbean) and national (Barbados and Dominica) climate and health sectors with respect to arbovirus and vector surveillance and control, and climate monitoring and forecasting. Information was gathered through face-to-face interviews with key stakeholders.

**Regional climate sector**

The Caribbean Institute of Meteorology and Hydrology (CIMH) is the technical arm of the Caribbean Meteorological Organization (CMO). In this role, it supports the National Meteorological and Hydrological Services (NMHSs) of the CMO’s 16 Member States on issues related to weather, water and climate. The ministers responsible for meteorology in the CMO states are represented on the Council of the CMO, providing guidance and direction for the region. In May 2017, the CIMH was designated as the World Meteorological Organization’s (WMO) Regional Climate Center (RCC) for the Caribbean, expanding their role in the region and the countries that they serve. The Applied Meteorology and Climatology Section of the CIMH forms the core of the RCC, that provide products and services for sectors and the public, to move from data to information to services. The work of the RCC is guided by the WMO’s Global Framework for Climate Services (GFCS). CIMH works with 6 key sectors: water, agriculture and food security, disaster risk management, health, energy, as well as tourism - an additional sector that is not a current focus of the GFCS. CIMH plays a key role in building the capacity of the NMHSs in the region to expand their portfolio of services from weather and aeronautical, to include products and services related to seasonal climate. As CIMH has increased their climate services portfolio, they have increased engagement with sectoral stakeholders, guided by an analysis of sectoral needs through social science research. The Regional Climate Centre is attracting funding from donors interested in supporting the region’s adaptation to climate risk.

CIMH is seeking to support the health sector to solve urgent climate-sensitive health problems. CIMH places most of their emphasis on the near future (seasonal and interannual climate variability) time scale as opposed to the long-term climate change time scale.

The CIMH, in association with its network of NMHSs, jointly produce operational climate monitoring and outlook information on the seasonal timescale. Twice a year, at the beginning of the region’s major Wet and Dry Seasons, the Institute convenes the Caribbean Climate Outlook Forum (CariCOF) to present the suite of seasonal climate forecasts and discuss their societal implications with key stakeholder groups. The seasonal climate information is routinely published on the Caribbean RCC website, and is distributed via email to regional meteorologists and climatologists, and in particular a wide cross-section of national and regional sectoral practitioners via targeted bulletins. The NMHSs are responsible for re-distributing the seasonal climate forecasts to, or downscaling them for, their national stakeholders. The CIMH also convenes regional and national level meetings related to the Sectoral EWISACTs (Early Warning Information System across Climate Timescales) with sectoral stakeholders.

**Barbados and Dominica national climate sectors**

In the Caribbean, the national meteorological and hydrological services (NMHS) have historically focused on aviation meteorology. Their focus has been more operational than research oriented. However, in the last 10 years, they have begun to develop climate services and seasonal forecasts with support from CIMH, and this is an area that they would like to grow and develop. Currently, the NMHS of Barbados and Dominica do not have a mandate to work on climate and health.

The Barbados NMHS produces weather forecasts three times each day for Barbados, St. Vincent, Dominica. They also produces seasonal rainfall and drought outlooks using 10 rainfall stations, with the temperature data from two of these stations are used to provide temperature outlooks. Apart from temperature and rainfall, these two stations also have a long history of other climate parameters such as relative humidity and windspeed. On Dominica, there are two long term weather stations located at the airports (Canefield and Douglas Charles). A network of hydrometerological stations is being installed to improve coverage across the island. There are also 10 Domex rain gages, through a project with Yale University. Each month the NMHS offices submit rainfall and temperature data from their long-term stations to CAROGEN, an in-house developed online platform at CIMH that supports the ingestion of climate data from Caribbean countries and territories into a regional database as well as the generation of seasonal climate information products, including seasonal climate forecasts for the region.

**Regional health sector:**

Health is a GFCS priority, and a priority for Caribbean societies that are impacted by climate sensitive diseases such as dengue, Zika and Chikunguya. The Caribbean Public Health Agency (CARPHA) was established in 2011 and became operational in January 2013. As the technical arm of CARICOM with responsibility for health, CARPHA has a mandate to protect the health, the environment and well-being in the Caribbean. Climate change and health (CCH) is an emerging area for CARPHA and is aligned to the WHO Strategy (CCH) and the priorities of the Strategic Roadmap for Climate Change and Health in the Caribbean. Six strategic imperatives were developed by the expert panel on climate change and health to build resilience in the health sector in the region.

CARPHA has twenty-six Member States that are supported to manage current and emerging public health issues. Guided by the Caribbean Cooperation in Health (CCH-IV) and the CARPHA Strategic Plan (2016 – 2025) assistance is provided to the Ministries of Health and other relevant Ministries to increase capacities to respond to outbreaks and to mobilize resources to develop and implement regional public goods. An early warning system for arboviruses aligns with CARPHA's mandate to assist Member States to reduce risk and manage arboviral diseases in the Caribbean.

The Pan American Health Organization (PAHO) is the specialized international health agency for the Americas and the Regional Office for the Americas of the World Health Organization (WHO). PAHO works with countries throughout the region to fight communicable and noncommunicable diseases and their determinants, to strengthen health systems and services, and to respond to emergencies and disasters. PAHO has 52 members countries and territories, under whose guidance it works to set regional health priorities and mobilize action to address health problems on a biannual basis.

The PAHO Strategic Plan (2014-2019) entitled “*Championing Health: Sustainable Development and Equity*”, sets out the strategic direction of the Organization based on the collective priorities of its member states and country focus. It also ensures programmatic alignment with the global health objectives of the WHO. Climate and health has been mandated as a high priority for PAHO, as it is a central part of the World Health Organization’s (WHO) global strategic plan. PAHO’s Climate Change and Health Program aims to prepare health systems through early warning, better planning and implementation of prevention and adaptation measures within the health sector and with other sectors

**Barbados health sector: disease surveillance and vector control**

The effects of climate on health is a relatively high priority in the Ministry of Health (MoH) in Barbados, but as it is a relatively new area, there is no mandate to do this work. Arboviruses are a high priority because of the high burden of disease. The vector control unit of the Environmental Health Department (EHD), in addition to routine monitoring of vectors and vector borne diseases across the island, is responsible for vector surveillance and control in ports of entry, and vector control preceding mass gathering events. The unit is also responsible for purchasing insecticides that are distributed to decentralized polyclinics located across the island. The mandate of the environmental health officers, who are located at the polyclinics, is to conduct routine vector control and surveillance, investigate foci of disease outbreaks and to visit each home at least two times per year.

The surveillance unit in the EHD is responsible for arbovirus disease surveillance. They receive data from hospitals and polyclinics. Physicians and nurses are required to fill out a paper notification form and send the form to the epidemiological unit, where the data are entered into a central database. The surveillance unit liaises with the Best-dos Santos Public Health laboratory (the national reference laboratory) to record laboratory confirmed cases. This is a passive surveillance system, and there is significant underreporting of suspected cases of dengue fever by private physicians. The surveillance unit compares weekly arbovirus cases from the past 5 years, where available, excluding outbreak periods, to current cases to determine if there is "above normal" transmission.

The national reference laboratory conducts dengue, chikungunya and Zika virus diagnostic testing for any individual with a suspected infection. Diagnostics include dengue virus IgM/IgG and chikungunya IgM/IgG using commercial ELISA kits. They also conduct real time PCR using the Centers for Disease Control and Prevention (CDC) Trioplex assay (since Sept 2016) for dengue, chikungunya and Zika viruses. Dengue positive specimens are serotyped. PCR is used for individuals within 4 days of infection, and IgM is used if the individual has been sick for more than 5 days. IgM is used as a serology test for anyone who is PCR negative. Diseases with similar clinical presentation as dengue fever include hantavirus and parvovirus B19.

Arbovirus case data have been georeferenced to the enumeration district level (census districts) since 2013. About 90% of suspected and confirmed cases are mapped, except when there is an outbreak, because the large number of cases exceeds the capacity to map them. Georeferencing is conducted by the national vector control unit of the MoH.

When a case is confirmed or suspected, the EHD officers at the polyclinics intervene quickly to prevent further transmission. Chemical larvicides used include temefos (5% granular form; main larvicide used), Aquatain (silicone based compound), Methoprene (juvenile growth inhibitor); and larvivorous fish in some cases. The adulticide used for fogging is 95% malathion with 5% diesel. *Aedes albopictus* has not been detected. One of the major issues related to vector control is the lack of regulation of private sector vector control companies hired by companies and high-income communities. The MoH does not receive information regarding the type of insecticides used, or how and where they are used, potentially increasing the risk of insecticide resistance.

**Dominica health sector: disease surveillance and vector control**

In Dominica, climate and health is a priority area, and arboviruses are important because of the high burden of disease. The health sector has put emphasis on mainstreaming climate change adaptation within its overall activities. The EHD has a very broad mandate, working in nine program areas including vector control, food safety, occupational health, port health, school health, institutional hygiene, among others. The EHD has the mandate to monitor and to ensure that the activities of the public do not have a negative impact on the environment, and that the environment does not have a negative impact on the health of the population.

Since 2006, the MoH has outsourced vector control and surveillance to the company National Pest. The EHD monitor and supervise National Pest by also conducting surveillance and control on a quarterly basis; they sample 30% of homes in a cross sectional and representative sample. *Aedes albopictus* has not been detected. Each home across the island is visited about four times per year by the EHD officer and National Pest. National Pest uses bacillus thuringiensis israelensis (BTI) for larviciding, due of temefos resistance, and they use permethrin and malathion for fogging during outbreaks and mass gatherings. They also use larvivorious fish as a form of biological control. They conduct vector surveillance in ports, and vector control when needed.

When a patient is diagnosed with dengue, a nurse at the clinic is responsible for filling out a paper form, and sending the form each week to the Health Information Unit (HIU). At the same time, the EHD officer in the district is notified, to ensure a rapid intervention. The EHD officer, community health nurse, and National Pest work together to intervene in all homes around the case within 200 meters. Since 2015, the EHD has georeferenced cases of arboviral disease.

Serum samples from suspected arbovirus cases are sent to CARPHA in Trinidad for laboratory confirmation, which often takes three weeks. There is no national reference laboratory in Dominica for diagnostics of arboviruses. Underreporting of dengue cases is a major problem, especially from the private sector.
