## Supplement 2 Text for "Co-developing climate services for public health: stakeholder needs and perceptions for the prevention and control of *Aedes*-transmitted diseases in the Caribbean"

**S2 Text. Interview and survey instruments**. This document contains (1) questions about climate information for arboviral control, used in interviews with climate and health decision makers, managers and expert practitioners, (2) interview questions regarding climate and health mandates, competencies, and data, (3) survey for health sector decision makers, managers, and expert practitioners.

**1. Climate and health interview for decision makers, managers, and expert practitioners**

**SECTION 1: BACKGROUND INFORMATION**

Professional Background:______________

What is the name of the institution where you work? ___________

What is your position at your institution?______________

Time working in the sector ________years ________months

Indicate the jurisdiction of your institution (check all that apply)

☐ Regional (Caribbean) ☐ National ☐ Community

What is the approximate number of technical staff in your department?

What is the principal function of your department/office in your institution? *Prompts: vector borne diseases, disease and vector control/surveillance, climate, climate change, climate-health linkages, climate-health communications, Disaster Risk Reduction (DRR).*

**SECTION 2. CLIMATE AND RELATED INFORMATION FOR VECTOR BORNE DISEASES**

In your experience, how does climate affect directly and indirectly the health of people in your jurisdiction? *Prompts: water-borne diseases, vector-borne diseases, respiratory diseases, heat waves, hurricanes, droughts, floods, fires, mental health (depression, anxiety), air quality, water quality, unsafe sewerage, food security*

In your jurisdiction, what are the key factors that increase the risk of epidemics of diseases transmitted by *Aedes aegypti* (dengue fever, chikungunya, and Zika fever)? *Prompts: human movement, insecticide resistance, lack of community engagement/mobilization, lack of public health education, emergence of a new virus, insufficient staff/resources for vector control and diagnostics, high social vulnerability (e.g., urban poverty), climate conditions, basic services to the communities (drinkable water, sewage treatment).* In your jurisdiction, what climate factors trigger epidemics of diseases transmitted by *Aedes aegypti* (dengue fever, chikungunya, and Zika fever)? *Prompts: rainfall, air temperature, relative humidity, El Niño indices (ENSO)*.*

**(ENSO) is a naturally occurring phenomenon that involves fluctuating ocean temperatures in the equatorial Pacific. When sea surface temperatures are warmer than average (+0.5 deg C from the long term average), this is referred to as an El Niño event. When sea surface temperatures are cooler than average (-0.5 deg C from the long term average), this is referred to as a La Niña event.*

Does your department/office use weather or climate information to plan or implement sector interventions that may benefit or lead to the reduction of VBDs? Or specifically for *Aedes aegypti*?

Do you know if there are early warning systems for *Aedes aegypti* transmitted diseases in your jurisdiction? Do these early warning systems use climate information?

Do you know if your National Climate Change Team/Committee or National Disaster Management Committee address public health measures related to vector borne diseases? Have you been directly involved?

**SECTION 3. CO-DEVELOPMENT OF CLIMATE SERVICES**

From what institutions and in what format do you receive climate information? *Prompts: Caribbean Institute of Meteorology and Hydrology (CIMH), National Met Service, other public institutions, NGOs, academia, private sector, technical advisors, program directors, WHO/PAHO/WMO, CARPHA, Scientific advisor, Scientific journals*

What would you like to see as climate services (final products) that result from a collaboration between your sector and the climate & health sector? How would you use these products in your daily work? *Prompts: Training workshops, interactive GIS platform online, interactive excel spreadsheets, Climate and health bulletins, climate and health forums (quarterly), annual climate-health regional meetings, internal meetings between directors (bimonthly, quarterly), website*

What factors **limit** or **enable** your department to work more closely with the climate & health sector?

- Modeling capacity: human capacity, software, hardware
- Trained personnel and technical capacity
- Evidence of knowledge about climate & vector borne diseases (VBDs) interactions
- Prior experience with early warning systems or other climate services.
- Availability of financial resources
- Efficiency in the management and distribution of financial resources
- Coordination for cross cutting interventions for VBDs management
- Mobilization and coordination with local communities (Community-Based interventions)
- Strong leadership (vision)
- Organizational structure

What can be done to make your sector more interested or engaged in climate services for health? *Prompts: More information and understanding (training), time, expert advisory (scientific, operative), funding, leadership, peer support academia collaboration partnerships with NGOs, private sector, community leaders*

What kind of activities or trainings would be most useful for your institution?

**SECTION 4. MULTI-SECTOR ENGAGEMENT FOR CLIMATE AND HEALTH (VBDs)**

What other sectors/organizations do you currently partner with on work related to VBDs? *Prompts: National Met Service, environment, risk management, education, NGOs, private sector, community-based organizations.*

What other sectors/organizations/stakeholders would you like to partner with to improve the control of *Aedes aegypti* transmitted diseases? *Prompts: National Met Service, civil society, private sector, NGOs, academia, community organizations.*

How does climate and health (vector proliferation) fit within their institutional priorities/mandates/competencies?

What strategies would improve the collaborations between institutions working in the area of climate and VBDs? *Prompts: Institutional Agreement (MOU); Climate, Risk and Health Forums; Specific protocols of collaboration; Climate and health bulletins; data sharing agreements*

**SECTION 5. NETWORKING**

Would you mind sharing the names of others who you think would be useful to interview on this subject? We are looking for other key stakeholders with some awareness or interest, and who may be able to increase their agency’s engagement. (Get contact information)

Would you be interested in receiving information from the results of this project by email?

Do you have any questions for me/us? Is there anything else you’d like to share?

You’ve been so helpful; I really appreciate the time you’ve taken to talk with me today. Do you mind if we contact you in the future with any follow-up questions that may emerge? Thank you very much.

**2. Interview: Climate and health mandates, competencies and data**

*Question for key informants in health and climate sectors.*

Please share an organizational diagram of the Ministry of Health of your country.

Please share links or copies of the laws and regulations that govern health care and the provision of health services related to vector-borne diseases in your country.

Please share links or copies of the laws and regulations that govern climate services for public health in your country.

Are there data sharing agreements between the public health sector and other agencies, such as the National Meteorological Services, Caribbean Institute of Meteorology and Hydrology, others?

**Disease case reports and surveillance**

1. What institution is responsible for disease surveillance?
2. How are cases of dengue fever, chikungunya and Zika reported (most likely passive surveillance?
3. How are cases diagnosed (clinical diagnosis, laboratory diagnosis)?
4. How are patients referred for laboratory testing? Do the patients come here or are blood/serum samples sent from the clinic?
5. Who conducts laboratory diagnoses of dengue/Zika/chikungunya cases?
6. What lab methods are used?
7. Is there historical data available on the prevalence of different DENV serotypes?
8. Is there targeted surveillance of ZIKV in pregnant women and congenital complications in infants?
9. Is there active surveillance of infections in the community?
10. Is there a community surveillance/detection system in place?
11. What is the spatial and temporal resolution of existing case records?
12. Are there electronic patient records or another electronic epidemiological database centralized for the country?
13. Are cases georeferenced? By whom?

**Vector surveillance and control**

1. What institution is responsible for vector surveillance and control?
2. How does the public health sector decide where to intervene for vector control and surveillance? Examples: Houses of suspected/confirmed cases, door to door vector control, high risk neighborhoods, requested by the communities
3. What are the most important factors that are considered when planning vector control/surveillance for the coming year? The following season? Weeks or days?
4. Is vector control and surveillance decentralized or centralized?
5. What vector surveillance tools are used? Larval surveys, pupal surveys, ovitraps, CDC light traps, prokopack backpack aspirators, other?
6. How frequently are homes visited for vector surveillance control?
7. Is there focal control (in and around presumed positive cases)?
8. What are the primary control methods for *Aedes aegypti*? Larvicide (temefos or BTI), indoor residual spraying (what chemical?), ultra low volume fogging (what chemical?), elimination of containers, use of petroleum products in standing water, biological control (copepods or larvivorous fish)?
9. Do you conduct surveillance of insecticide resistance? If yes, when? Where? How often?
10. Is there a national electronic database with historical vector surveillance data? What spatial and temporal resolution?
11. Has the vector surveillance data been georeferenced? By whom?
12. What is the timing (schedule) of vector control interventions (e.g., seasonal, monthly, random)?
13. Surveillance of *Aedes aegypti*, includes which of the following variables and what is the frequency (weekly, monthly, seasonal, random) of the data collected:

- GPS points (geographic location)
- Specific Address
- Elevation
- Local precipitation
- Local Temperature
- Breteau Index
- House Index
- Pupal indices
- Container indices
- Adult indices
- Housing characteristics
- Presence of febrile infections
- Household demographics
- Insecticide resistance
- Indoor versus outdoor location of adult mosquitoes

**National census data**

1. When was the last national household census conducted?
2. Are the census records geoferenced? At what spatial resolution?
3. Have vulnerability maps been created to identify the characteristics of high-risk communities (using census data)?

**Climate data (CIMH/National Meteorology Service)**

1. Who generates weather forecasts, seasonal forecasts, climate change projections?
2. How does your institution distribute their climate information?
3. How do CIMH and the NMS interact to support the forecast operations of the Caribbean Outlook Forum?
4. Is there a standardized database?
5. Spatial and temporal scale of meteorological data?
6. Location of met stations?
7. Automatic/manual stations?
8. What are the barriers/limitations and resources available for forecasting?

**Climate and health capacity for supporting an early warning system:**

1. Current resources and needed resources: human personnel (modelers), soft/hardware.
2. Who are the key technical personnel involved in future capacity building for the development and implementation of an EWS?

**3. Climate and health survey for public health decision makers, managers, and expert practitioners**

**SECTION 1: BACKGROUND INFORMATION**

Country:__________

Institution:__________

Department/unit:___________

What is your position at your institution?______________

Time working in the sector ________years ________months

Indicate the jurisdiction of your institution (check all that apply). Your jurisdiction refers to the geographic area (e.g., country, district, parish) which your department and institution serves.

☐ Regional (Caribbean) ☐ National ☐ Community ☐ Other:___________

Age: ☐ 18-30 ☐ 30-40 ☐ 40-50 ☐ 50-65 ☐ >65 ☐ Prefer not to reply

Gender: ☐ Male ☐ Female ☐ Other ☐ Prefer not to reply

Highest level of education completed:

☐ Primary education ☐ High School ☐ Bachelor’s Degree ☐ Master’s Degree ☐ Ph. D. ☐ Other:____

Professional Background:

☐ Public health

☐ Medical

☐ Entomology

☐ Risk

☐ Management/administration

☐ Environment

☐ Engineering

☐ Social Work

☐Other:________________

**SECTION 2. CLIMATE INFORMATION FOR VECTOR BORNE DISEASES**

In your experience, which of the following factors are important in triggering epidemics of diseases transmitted by *Aedes aegypti* (dengue fever, chikungunya, and Zika fever) in your jurisdiction?

| **Risk factors** | **Not important** | **Slightly important** | **Moderately important** | **Important** | **Very important** | **I don’t know** |
| --- | --- | --- | --- | --- | --- | --- |
| Introduction of a new virus to a susceptible population |  |  |  |  |  |  |
| Mosquitoes that are resistant to insecticides |  |  |  |  |  |  |
| Limited community engagement/mobilization |  |  |  |  |  |  |
| Lack of community knowledge and awareness |  |  |  |  |  |  |
| Insufficient staff/resources for vector control |  |  |  |  |  |  |
| High-risk housing conditions |  |  |  |  |  |  |
| Human movement |  |  |  |  |  |  |
| Heavy rainfall |  |  |  |  |  |  |
| Drought conditions |  |  |  |  |  |  |
| Warmer air temperatures |  |  |  |  |  |  |
| El Niño or La Niña events* |  |  |  |  |  |  |
| Water storage behavior |  |  |  |  |  |  |
| Economic barriers to mosquito control by households (e.g., cost of screens or insecticide) |  |  |  |  |  |  |
| Low risk perception by communities |  |  |  |  |  |  |

**The El Niño Southern Oscillation (ENSO) is a naturally occurring phenomenon that involves ups in downs in ocean temperatures for 6 to 18 months in the east of the equatorial Pacific. When those temperatures are significantly warmer than average, this is referred to as an El Niño event, which in much of the Caribbean is linked to droughts. When the temperatures are significantly cooler than average, this is referred to as a La Niña event, which is linked to heavy rainfall and flooding*

Please describe any other risk factors not included in the prior question:____________________________

Have you received information on the effects of climate on vector-borne diseases?

☐ Yes ☐ No ☐ I don’t know

Does your health department use climate information to plan or implement disease and vector control interventions? ☐ Yes ☐ No ☐ I don’t know

If your answer is YES, what is the source of the climate information (mark all that apply and specify)?

☐ Caribbean Institute of Meteorology and Hydrology (CIMH) ☐ National Meteorological Service: _____

☐ Universities ☐ Private institutions ☐ Others_______

If you had access to additional climate information, (1) what information would be helpful and (2) how could you use this information in your daily work to reduce the risk of vector-borne diseases?

**SECTION 3: CLIMATE PERCEPTIONS AND PUBLIC HEALTH RESPONSES**

**“Climate variability”** refers to fluctuations in climate around the long-term average, which occur over months, to seasons, to years. El Niño and La Niña are good examples of features of climate variability that affect weather and our society in the Caribbean. Examples of impacts of climate variability include long-term flooding and recurrent flash floods, droughts, heat waves, amongst others.

How much do you agree or disagree with the following statements?

|  | 1  Strongly Disagree | 2  Disagree | 3  Neither agree nor disagree | 4  Agree | 5  Strongly Agree | 6  Don’t know |
| --- | --- | --- | --- | --- | --- | --- |
| My jurisdiction is currently experiencing one or more serious public health problems as a result of climate variability. |  |  |  |  |  |  |
| My jurisdiction is currently experiencing an increased risk of diseases transmitted by *Aedes aegypti* due to climate variability. |  |  |  |  |  |  |
| In the next 20 years, my jurisdiction will experience increasing risk of diseases transmitted by *Aedes aegypti* due to climate variability. |  |  |  |  |  |  |
| I am worried about the impact of climate variability on the health and well-being of people in my jurisdiction. |  |  |  |  |  |  |
| The effects of climate variability on the health of people in my jurisdiction is an urgent problem. |  |  |  |  |  |  |
| There are options/solutions to reduce the effects of climate variability and to improve the health of people in my jurisdiction. |  |  |  |  |  |  |
| The people in my jurisdiction are worried about the effects of climate variability on their health and well-being. |  |  |  |  |  |  |
| My health department currently has ample expertise to assess the potential public health impacts associated with climate variability that could occur in my jurisdiction. |  |  |  |  |  |  |
| Dealing with the public health effects of climate variability is an important priority for my health department. |  |  |  |  |  |  |
| I am knowledgeable about the potential public health impacts of climate variability. |  |  |  |  |  |  |
| The other relevant senior managers in my health department are knowledgeable about the potential public health impacts of climate variability. |  |  |  |  |  |  |
| My health department currently has ample expertise to create an effective plan to protect local residents from the health impacts of climate variability. |  |  |  |  |  |  |
| My health department currently has sufficient resources to effectively protect local residents from the health impacts of climate variability. |  |  |  |  |  |  |
| My health department is able to effectively communicate the health impacts of climate variability to local communities. |  |  |  |  |  |  |

**SECTION 4: EARLY WARNING SYSTEMS FOR VECTOR-BORNE DISEASES**

Do you know what is an early warning system for disease epidemics? ☐ Yes ☐No ☐ I don’t know

Do you know if there are early warning systems for *Aedes aegypti* transmitted diseases in your jurisdiction? ☐ Yes ☐No ☐ I don’t know

Do these early warning systems use climate information? ☐ Yes ☐No ☐ I don’t know

*In your words*, describe an optimal early warning system to predict and prevent local epidemics of mosquito-borne diseases, like dengue fever: ______________________________________________________________________________

______________________________________________________________________________

If there was an early warning system created to predict epidemics of dengue fever, chikungunya and Zika fever, how would you prefer to receive the warnings or alerts (mark all that apply)?

☐ Climate and health bulletins (PDF) by email

☐ Climate and health forums (quarterly)

☐ Annual climate-health regional meetings

☐ Internal meetings within your department

☐ Interactive GIS platform online

☐ Interactive excel spreadsheets

What are the strengths of your health department, **which would enable** the implementation an early warning system for *Aedes aegypti* transmitted diseases? (mark all that apply)

☐ Computer programming expertise

☐ Geographic information system (GIS) expertise

☐ Statistical and/or modeling expertise

☐ Computing hardware to operate the early warning system (computers, data servers)

☐ General knowledge of climate and vector borne diseases

☐ Organizational structure

☐ Prior experience with early warning systems or other climate services

☐ Availability of financial resources

☐ Efficiency in the management and distribution of financial resources

☐ Effective vector and disease surveillance infrastructure

☐ Effective public health messaging/education

☐ Mobilization and coordination with local communities

☐ Strong coordination with other institutions/NGOs/private sector

☐ Strong leadership

Which areas would **need to be strengthened** in order to implement an early warning system for *Aedes aegypti* transmitted diseases? (mark all that apply)

☐ Computer programming expertise

☐ Geographic information system (GIS) expertise

☐ Statistical and/or modeling expertise

☐ Computing hardware to operate the early warning system (computers, data servers)

☐ General knowledge

☐ Organizational structure

☐ Prior experience with early warning systems or other climate services

☐ Availability of financial resources

☐ Efficiency in the management and distribution of financial resources

☐ Effective vector and disease surveillance infrastructure

☐ Effective public health messaging/education

☐ Mobilization and coordination with local communities

☐ Strong coordination with other institutions/NGOs/private sector

☐ Strong leadership

**SECTION 5. CAPACITY BUILDING**

In your department, are there people with expertise in the following GIS, statistical, programming, or database software? (mark all that apply)

☐ ArcGIS

☐ QGIS

☐ Tableau

☐ Epi Info

☐ R

☐ SPSS

☐ SAS

☐ Google Earth

☐ Microsoft Excel

☐ Microsoft Access

☐ SQL

☐ Visual Basic

☐ Java

☐ Other:___

What kind of activities or trainings would be useful for your institution? (mark all that apply)

☐ Climate forums: information and data for seasonal forecasts

☐ Climate forums: data and models for 5 years – scenarios, outlooks, and other tools.

☐ Technical workshop on how to use climate information to predict epidemics)

☐ Use of GIS (digital maps) to identify areas at risk of vector-borne diseases.

☐ Time series analysis of health and climate data

☐ How to communicate the effects of climate on health to local communities

☐ Simulations of an early warning system for epidemics

☐ Other:__________________________________________________

Thank you for taking the time to complete this survey!

Would you like to receive a summary report of the results? ☐ No ☐ Yes. Email:_____________________
