## Supplement 3 Text for "Co-developing climate services for public health: stakeholder needs and perceptions for the prevention and control of *Aedes*-transmitted diseases in the Caribbean"

**S3 Text. Forecast scenarios discussed in the Barbados stakeholder workshop.** This activity was conducted at a national consultation at the PAHO in Bridgetown, Barbados, in April 2017, with 27 representatives from the national Ministry of Health (MoH) of Barbados, the Barbados Meteorological Services, the CIMH, and the PAHO. Participants were divided into small groups that included representatives from climate and health sectors. Groups were asked to respond to different forecast scenarios (2 week, 3 month, and 1 year forecasts of *Aedes aegypti* larval indices and dengue incidence). They were asked to identify the actions that they would take in response to alerts at each time scale, and they discussed the utility of a vector versus disease forecast. Results were identified by coding the transcripts of audio recordings.

**Activity instructions:**

- Break into 3 groups (will count off). In each group, one representative of CIMH and the national meteorological service.
- Assign a note taker, a moderator, and a presenter.
- 30 minutes to discuss in small groups. Record key points on flipchart paper.
- 30 minutes (7-10 min per group) to present and discuss with everyone.

**What actions would your sector take in response to the following alerts?**

| **Short term (2 week forecast):** | **Results** |
| --- | --- |
| - In two weeks there is a high probability that *Aedes aegypti* larval indices will increase. - In two weeks, there is a high probability of a dengue outbreak. | - At this time scale, the health sector would use the same strategies as always, but would increase education and mobilization, especially in the known hot spots - Short-term forecasts are important. They provide some lead-time to respond to a dengue outbreak. - Informants engaged in vector control indicated that alerts even at 2 weeks would be sufficient to allow them to implement source reduction programs in high risk areas and to prepare the population. |
| **Medium term (3 month forecast):** |  |
| - In the next three months (May-July), there is a high probability that *Aedes aegypti* larval indices will increase. - In the next three months (May-July), there is a high probability of a dengue outbreak. | - At this time scale, the health sector would be better able to plan with stakeholders, mobilize the field team, look at trends, and create bulletins for community mobilization. - Quote: “a year can feel like a long time away. With 3 months, there will be a sense of urgency and you can do meaningful activities, although there might not be new resources”. - Informants from the national reference laboratory, responsible for arbovirus diagnostics, indicated that they needed at least 6 months of lead-time in order to effectively procure diagnostic reagents in Barbados. |
| **Long term (1 year forecast):** |  |
| - Next year, during May to July, 2018, there is a high probability that *Aedes aegypti* larval indices will increase. - Next year, during May to July, 2018, there is a high probability of a dengue outbreak. | - At this time scale the health sector could better lobby for the needed financial support, allowing for more effective budgeting. They would also be able to better mobilize and train community members. They would able to monitor and evaluate interventions, and conduct a needs assessment to inform plans. - They suggest focusing on engaging critical stakeholders, such as policy makers and leaders in the MoH and Minister of Finance. |
