## Supplement 4 Table for "Co-developing climate services for public health: stakeholder needs and perceptions for the prevention and control of *Aedes*-transmitted diseases in the Caribbean"

**S4 Table. Types of software currently used in health departments.** Results from surveys are shown as % (n).

| **Categories** | **% (n)** |
| --- | --- |
| Microsoft Excel | 75.0% (24) |
| Epi Info | 62.5% (20) |
| Google Earth | 56.3% (18) |
| Microsoft Access | 50.0% (16) |
| ArcGIS | 28.1% (9) |
| QGIS | 21.9% (7) |
| SPSS | 21.9% (7) |
| Java | 9.4% (3) |
| Visual Basic | 3.1% (1) |
| Stata | 3.1% (1) |
| SQL | 0 |
| R | 0 |
| Tableau | 0 |
| SAS | 0 |
