## Supplement 5 Table for "Co-developing climate services for public health: stakeholder needs and perceptions for the prevention and control of *Aedes*-transmitted diseases in the Caribbean"

**S5 Table. Preferred training activities identified by health sector survey respondents.** Results from surveys are shown as % (n).

| **Categories** | **% (n)** |
| --- | --- |
| Technical workshop on how to use climate information, data & models and other tools to predict epidemics) | 78.1% (25) |
| Use of GIS (digital maps) to identify areas at risk of vector borne diseases. | 75.0% (24) |
| How to communicate the effects of climate on health to local communities | 75.0% (24) |
| No response | 6.3% (2) |
