## Supplement 6 Table for "Co-developing climate services for public health: stakeholder needs and perceptions for the prevention and control of *Aedes*-transmitted diseases in the Caribbean"

| **S6 Table. Current use of climate information and early warning systems reported by survey respondents.** Results shown as % (n). | | | | |
| --- | --- | --- | --- | --- |
| **Questions** | **No response** | **Don't know** | **No** | **Yes** |
| Have you received information on the effects of climate on vector-borne diseases? | 0 (0) | 0 (0) | 31.3 (10) | 68.8 (22) |
| Does your health department use climate information to plan or implement disease and vector control interventions? | 0 (0) | 18.8 (6) | 31.3 (10) | 50 (16) |
| Do you know what an early warning system for disease epidemics is? | 3.1 (1) | 0 (0) | 18.8 (6) | 78.1 (25) |
| Are there early warning systems for *Aedes aegypti* transmitted diseases in your jurisdiction? | 6.3 (2) | 12.5 (4) | 40.6 (13) | 40.6 (13) |
| Do these early warning systems use climate information? | 28.1 (9) | 37.5 (12) | 21.9 (7) | 12.5 (4) |
