## Supplement 7 Table for "Co-developing climate services for public health: stakeholder needs and perceptions for the prevention and control of *Aedes*-transmitted diseases in the Caribbean"

**S7 Table. Preferred way of receiving information from an early warning system that predicts arbovirus epidemics.** Results shown as % (n).

| **Categories** | **% (n)** |
| --- | --- |
| Climate and health bulletins (PDF) by email | 90.6% (29) |
| Online interactive GIS platform | 65.6% (21) |
| Internal meetings within your department | 59.4% (19) |
| Climate and health forums (quarterly) | 34.4% (11) |
| Annual climate-health regional meetings | 25.0% (8) |
| Interactive excel spreadsheets | 25.0% (8) |
